# The Neurophysiological Modality Effect in Native and Second Language Processing: An ERP Study

**DOI:** 10.1101/2022.12.17.520859

**Authors:** Daniel Gallagher, Masataka Yano, Shinri Ohta

## Abstract

Experimental psychology has long discussed the modality effect, whereby the stimulus modality significantly affects retention of the information presented. In neurolinguistics, however, the effect of stimulus modality on language processing has gained little attention. We conducted a multi-modal event-related potential (ERP) experiment on both native and non-native Spanish speakers to investigate the possibility of a neurophysiological modality effect in language processing. Using morphosyntactically violated and orthographically/phonologically violated stimuli, we elicited a robust N400 and P600 in native speakers. We showed that the N400 has consistent features across modalities, while the P600 has modality-specific features. Specifically, the auditory evoked P600 was characterized by a more gradual slope and a later peak than the visual evoked P600. We discuss this in detail along with other modality effects observed post sensory perception. Among second language (L2) learner groups, those with higher proficiency exhibited more nativelike neurophysiological responses in both modalities when compared to those with lower proficiency. We additionally observed fewer modality-specific differences in low proficiency learners than in higher proficiency learners suggesting that modality-specific specialization in language processing comes with increased proficiency. We further discuss the question of modality-specific differences in the process of neurophysiological nativization, whereby L2 learners’ ERPs become increasingly on nativelike.

## 1 Introduction

The modality effect, whereby the stimulus modality significantly affects behavioral performance, is well-known in experimental psychology. The modality effect initially referred to greater recall of auditory stimuli in short-term memory tasks. However, depending on the characteristics and recency of the stimulus, a superior recall could also be produced in the visual modality (Engle & Mobley, 1976; Penney, 1989; Watkins, 1972). More recently, the modality effect has come to be commonly associated with increased retention of information that is simultaneously presented graphically and auditorily, as opposed to graphically and textually, and meta-analyses have shown this to be a reliable effect to varying degrees (Ginns, 2005; Reinwein, 2012). Great efforts have been dedicated to modeling and explaining the neurophysiological cause of the effect in terms of working memory. Baddeley’s model of verbal short-term memory (A. Baddeley, 1992), Penney’s separate streams hypothesis (Penney, 1989), and Sweller’s cognitive load theory (Sweller, 1988) have been experimentally investigated for decades and are still widely influential in psychological and pedagogical studies. Meanwhile, in neurolinguistics, the effects of stimulus modality are rarely given their due attention and are often outright ignored. The tacit assumption is that language processing is modality independent, and that, beyond very early stages, language processing does not differ much between reading and listening. But, to what extent is this true?

In this study, we use a multi-modal event-related potential (ERP) experimental paradigm to address this questions and to investigate the existence of a neurophysiological modality effect in native-language (L1) processing and second language (L2) acquisition. Contingent upon its existence, we further set out to characterize the latency and distribution of the ERP components associated with the modality effect.

### 1.1 The Basis for a Language Processing Modality Effect

As a simple matter of logical deduction, the functional anatomical pathways for sensory perception alone necessitate modality-specific processing. Audition begins with sound waves entering the ear canals. Sound vibrations are converted at the cochlea to an electrical signal, which is sent to the medial geniculate complex in the thalamus, where the signal is in turn relayed to the primary auditory cortex in the temporal lobe (Lee, 2015). From here, depending on the nature of the sound, the signal projects onto many different cortical areas. Vision, by contrast, begins with light entering the eye, where the retina converts it to an electrical signal. This signal is relayed via the lateral geniculate nucleus in the thalamus to the primary visual cortex in the occipital lobe, from where the signal is mapped onto many different cortical areas (Stoerig, 2001).

Up to this point, the two processing streams are sensory-perceptual in nature, not linguistic. Hereafter, acoustic signals are mapped onto articulatory representations and meaning (Hickok & Poeppel, 2007), while orthographic input is relayed to the visual word form area, where its orthographic representation is mapped onto its phonological representation and/or meaning (Price, 2012). Furthermore, functional differences show that visual word recognition unfolds by both a feedforward system, in which phonology mediates lexical access, and a feedback system, in which the semantic representation interacts directly with orthography perception (Carreiras, Armstrong, Perea, & Frost, 2014; Wheat, Cornelissen, Frost, & Hansen, 2010). In contrast, auditory word recognition has a more direct path for lexical access whereby phonology is directly mapped onto semantic representations. This removes the need to store orthographic representations in working memory, ultimately placing less demand on verbal working memory (Crottaz-Herbette, Anagnoson, & Menon, 2004). Thus, beyond simple sensory-perceptual processes, there at least exist functional anatomically based modality-specific differences in early language processing even post sensory perception.

Aside from the aforementioned functional neuroanatomical features of word recognition, three particular arguments can be made to suggest the likelihood of a neurophysiological modality effect in language processing. The first pertains to the temporal features of processing auditory stimuli compared to visual stimuli; the second appeals to the evolution of spoken and written language; the third argument springs from well-known effects related modality-specific language perception.

Regarding temporal features of stimulus processing, auditory stimuli elicit a reaction time faster than visual stimuli by about 50 ms (Shelton & Kumar, 2010). Similarly, auditory perception has higher temporal resolution than visual perception (Stauffer, Haldemann, Troche, & Rammsayer, 2012). These phenomena seem to favor the auditory modality, but the physical features of auditory and visual stimuli differ in such a way that may favor visual language processing. Namely, auditory information unfolds in a temporal, moment-to-moment order, whereas visual information becomes instantaneously available at the moment of presentation. Indeed, evidence based on the processing of transposed-letter nonwords (e.g., “cholocate” instead of “chocolate”) indicates that “good enough” orthographic representations of individual words are processed in much the same way as their correctly spelled versions (Lupker, Perea, & Davis, 2008; Meade, Grainger, & Holcomb, 2022). This suggests that visual words are processed holistically, at least to some extent, which, due to stimulus features, is not physically possible for auditory word processing.

An evolutionary argument can also be made for the existence of modality-specific differences in language-processing. Notably, humans have used spoken language for as long as 200,000 years (Pagel, 2017), while written language has only been used for a comparably short 5,000 years (Woods, 2011), meaning that in all likelihood, reading is accomplished by the repurposing (or exaptation) of a brain area that was originally dedicated to another task, i.e., the neuronal recycling hypothesis of reading (Dehaene & Cohen, 2007). By extension, reading disorders such as dyslexia, whose population prevalence is estimated at 10–15% (Fletcher, Lyon, Fuchs, & Barnes, 2007), outpacing analogous listening disorders like central auditory processing disorder, whose population prevalence is estimated at 2–3% (Chermak, 2001), may indicate an inefficiency in the neuronal recycling of exapted neural areas (Dehaene & Cohen, 2007).

Finally, when it comes to simultaneous exposure to both visual and auditory language cues, we can consider two well-known effects: The Colavita effect, and the McGurk effect. The Colavita effect results in adults (not children) responding only to visual stimuli when conflicting bimodal stimuli are simultaneously presented (Hirst, Cragg, & Allen, 2018). There is also the more well-known McGurk effect, whereby a video of lips producing /ga/ dubbed over with an audio recording of /ba/ results in participants unconsciously reconciling the two conflicting stimuli by interpreting them as a fusion of the two sounds, /da/ (McGurk & MacDonald, 1976). While the Colavita effect seems indicative of age-related increased reliance on visual input, the McGurk effect shows a reliance on sensory information from both modalities, wherever available. Thus, here too, we see that language processing can be greatly influenced by the modality of the stimulus.

While these three cases already offer convincing evidence for the possibility of a neurophysiological modality effect in language processing, we can also consider the robustly elicited non-linguistic modality-specific mismatch negativity (MMN) ERP component. Among the most prominently studied ERP components with well-known modality-specific features, the auditory MMN and its visual counterpart, the vMMN, index change-detection in response to deviance from an established repeating pattern, such as adjusting the frequency, duration, etc. of otherwise regularly-spaced tones in the auditory modality (see, e.g., Pakarinen, Takegata, Rinne, Huotilainen, & Näätänen, 2007) or by changing the presented letter in otherwise regularly-spaced presentations of a single letter (see, e.g., Kimura, 2012). Incredibly robust, the MMN has wide-ranging application, even as a prognostic measure in predicting positive outcomes for comatose patients (Luauté et al., 2005). The main difference between the auditory and visual counterparts of the MMN is the scalp distribution, with the vMMN having a parietooccipital distribution as opposed to the auditory MMN’s more fronto-central distribution (Näätänen et al., 2012; Wiens, Szychowska, & Nilsson, 2016). Importantly, the modality-specificity of the MMN demonstrates that similar processes can happen across modalities without a single amodal process accounting for all observed data. At the same time, fairly consistent onset latencies at around 200 ms post-stimulus onset, depending on task complexity (Kojouharova, File, Sulykos, & Czigler, 2019; Wiens et al., 2016), illustrate that separate processing streams can be close to timing-consistent across modalities.

Taken together, the functional neuroanatomy necessitates modality-specific language processing in the early stages. Whether or not such modality-specific differences exist in the later stages of language processing (e.g., during morphosyntactic processing) has not yet been thoroughly investigated, but the aforementioned evidence suggests a strong likelihood that an important part of language processing does not fit nicely into a single amodal language processing stream.

Previous studies directly investigating modality-specific differences in native language processing have largely targeted the N400 and P600 and concluded that language processing is mostly similar across modalities with only minor differences (Holcomb, Coffey, & Neville, 1992; Osterhout & Holcomb, 1993) or no differences at all (Balconi & Pozzoli, 2005). Other studies have referred to both similarities and differences across modalities only in passing (e.g., Osterhout, 1997; Van Berkum, Koornneef, Otten, & Nieuwland, 2007). Our findings for native speakers will be compared with these studies in the Section 4.1.

On the other hand, to our knowledge, it has never been investigated whether a modality effect exists in second language acquisition. But since non-native language processing differs from native language processing at lower proficiencies and then eventually converges to nativelike language processing (Steinhauer, 2014), even if no electrophysiological modality effect exists in native speakers, such an effect could exist in lower-proficiency L2 learners. Furthermore, it is conceivable that either modality becomes nativelike at an earlier stage of proficiency than the other modality.

### 1.2 Language Processing from the Perspective of Stimulus Modality

Electroencephalography (EEG) has become a staple method in neurolinguistics due to its being non-invasive and giving high temporal resolution. EEG signals time-locked to the onset of a stimulus, called ERPs, reveal the timing, magnitude, and, to a lesser extent, the distribution of electrical activity associated with specific neural processes. By looking at a specific ERP component, for example, the P600—a positive deflection over centro-parietal electrodes robustly elicited by grammatical violations (Osterhout & Holcomb, 1992)—and comparing these characteristics, it can be determined whether or not the modality effect manifests at the neurophysiological level during language processing.

In this study, we first consider the native speaker. Modeling language processing streams from the perspective of stimulus modality, we can conceive of five logically plausible models, as shown below in Figure 1.

**Figure 1.**
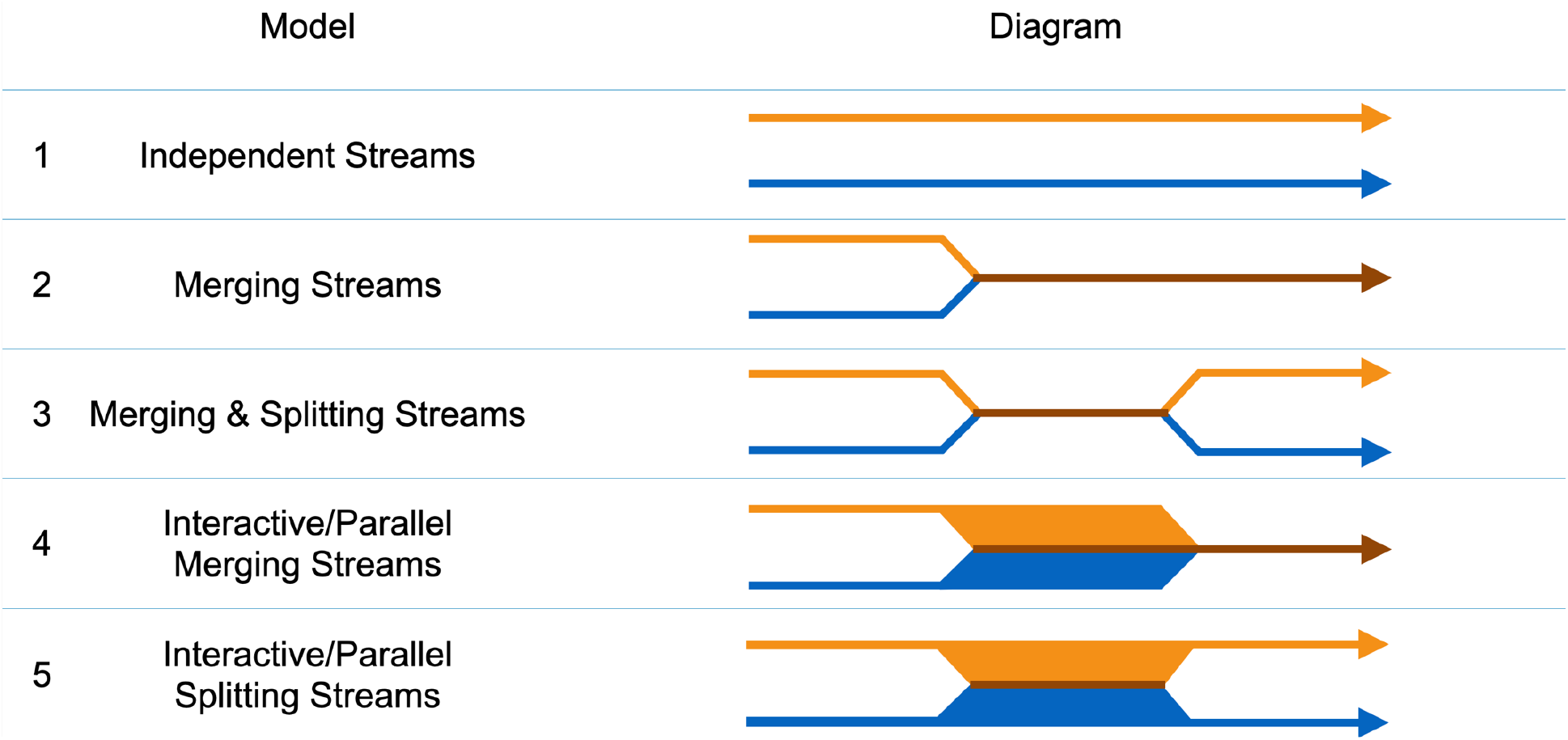
Logically conceivable models of language processing streams from the perspective of stimulus modality. Orange represents auditory-specific processing, blue represents visual-specific processing, brown represents modality-independent processing.

In Model 1, the auditory and the visual language processing streams are wholly independent and never merge or overlap. In Models 2 and 3, the processing streams are initially independent, but merge and either remain modality-independent (Model 2), or split back into modality-dependent processing streams (Model 3). In Models 4 and 5, the visual and auditory processing streams interact with or occur in parallel with a modality-independent language processing stream, where modality-specific processing either ceases in favor of modality-independent processing (Model 4) or outlasts modality-independent processing (Model 5). Below, we discuss the various models in Figure 1 in light of existing evidence.

However unlikely it may be that the brain has entirely separate processing streams for reading and listening, Model 1 is still a logically possible model that warrants consideration. While some neurological disorders or brain lesions may differentially affect reading (e.g., dyslexia) and listening (e.g., central auditory processing disorder), some types of aphasia such as primary progressive aphasia affect all domains of linguistic function without necessarily affecting other cognitive function (Gorno-Tempini et al., 2011). This alone provides compelling evidence for the existence of modality-independent linguistic mechanisms. But when considering the overwhelming volume of literature showing qualitatively similar ERP components, like the N400 and P600, in both modalities (see, e.g., Federmeier & Kutas, 1999; Federmeier, McLennan, de Ochoa, & Kutas, 2002; Hagoort & Brown, 2000; Kutas, Neville, & Holcomb, 1987), as well as the MEG and fMRI evidence pointing to shared cortical-activation sites, like the left inferior frontal gyrus and middle temporal gyrus (see, e.g., Bemis & Pylkkänen, 2013; Buchweitz, Mason, Tomitch, & Just, 2009; Michael, Keller, Carpenter, & Just, 2001), it is dubious at best to suggest a language processing model with no shared, amodal mechanisms. Thus, it is safe to eliminate Model 1 from our list of contenders, and conclude that, at some point, the two language processing streams must merge into or interact with/occur parallel with an amodal language processing stream. Nevertheless, if Model 1 were to be the true model, we would expect to see entirely different ERPs in the two modalities.

Model 2 may be how many people tacitly assume language processing works. If this model were to be correct, we would expect to see modality-specific processes (potentially, though not necessarily, beyond sensory-perceptual processes) up until a certain time, after which, the processing of auditory and visual stimuli is indistinguishable from one another.

Model 3 may initially seem counter-intuitive insofar as it is not apparent why the processing streams would need to split again after merging. However, one interpretation of the N400 and P600 is that they index prediction error backpropagation (Fitz & Chang, 2019). With this interpretation, it is reasonable to consider such errors propagating back to the level of representation, whether orthographic or phonological. This would cause a split in the merged processing stream, which would manifest as a late latency modality-effect. On the other hand, it is conceivable that this error backpropagation does not propagate all the way back to the level of sensory representation. Thus, a lack of a late-stage modality effect would only rule out the possibility of more extensive backpropagation, without ruling out backpropagation altogether. Furthermore, the presence of a late modality effect would of course not prove the backpropagation interpretation since other interpretations could still be plausible.

Model 4 differs from preceding models in that some portion of the processing stream involves modality-dependent processes interacting or occurring in parallel with modality-independent processes. For example, amodal grammar processing could theoretically rely on a double-checking procedure whereby a modality-specific stimulus (e.g., a pronoun) is stored in working memory and accessed again at the time of exposure to a syntactically related stimulus (e.g., a conjugated verb). If this model were to be true, we would expect to find some degree of overlap in ERPs (e.g., consistent P600 components), while still having a modality effect manifest in other ERP components. This model predicts that the final segment of language processing is amodal.

Model 5 is equivalent to Model 4 with the exception that the modality-independent processing ceases prior to modality-specific processing.

The specific nature of any modality effect discovered herein can shed light on which of the aforementioned models most accurately describes language processing. We therefore interpret the results of these experiments in light of these models and discuss the ramifications of our conclusions on neurolinguistics more broadly.

### 1.3 Expected Results: Morphosyntactic Violations

#### L1 Experiment

In order to make the tasks accessible to non-native speakers as well, the grammatical judgment task was very easy for native speakers. Given that, we expect native speakers to perform at a near-perfect level irrespective of stimulus modality. Thus, a ceiling effect is likely to prevent the observation of any behavioral modality effect.

As for the ERP results, morphosyntactic violations reliably elicit a P600 (Mancini, Molinaro, Rizzi, & Carreiras, 2011a). Since the discovery of the P600 (Osterhout & Holcomb, 1992), what exactly the P600 indexes has remained a hotly debated topic in neurolinguistics (see, e.g., Coulson, King, & Kutas, 1998; Frisch, Kotz, Yves con Cramon, & Friederici, 2003; Osterhout, McKinnon, Bersick, & Corey, 1996; Sassenhagen & Fiebach, 2019). As this is not a primary interest of the present study, it is sufficient for our purposes to simply note that a long-lasting relative positivity over centro-parietal electrodes (i.e., the P600) is expected in the morphosyntactically violated condition for both modalities. If the P600 is modality-independent, we expect there to be no differences between modalities. If the P600 is even partly composed of modality-dependent signals, we should expect a discrepancy in latency, amplitude, and/or distribution between the two modalities.

Aside from the P600, if a heretofore undescribed modality effect exists, its distribution may coincide with modality-specific neuroanatomy. Namely, the distribution of a neurophysiological modality effect may be temporal in the auditory modality (i.e., aligning with the primary auditory cortex) or occipital in the visual modality (i.e., aligning primary visual cortex). Although functional neuroanatomy would reasonably allow for such distributions, it does not preclude the possibility of other distributions, especially considering the poor spatial resolution of EEG data.

Additionally, given that the rules governing grammar are fundamentally not related to sensory modality, it is reasonable to expect that fewer neurophysiological modality effects (if any) would arise in morphosyntactic violation processing than in orthographic/phonological violation processing, which is inherently linked to stimulus modality.

#### L2 Experiment

Given that L2 participants have a variety of proficiency levels, and some may find the grammatical judgment task challenging, if there is a consistent behavioral modality effect, it may present itself in this task. Given that foreign language education in Japan often focuses on test preparation over more natural language use (Yoshida, 2003), a stronger performance with visual stimuli would be unsurprising.

L2 ERPs elicited by grammatical violations have been shown to evolve with the learner’s proficiency, whereby lower-proficiency learners responded to grammatical violations with N400s while higher-proficiency learners responded with P600s (McLaughlin et al., 2010). On the other hand, an L2 learner’s ERP is strongly affected by L1 transfer, or the extent to which L1 processing strategies are transferred over into L2 processing (for example, by the presence of similar grammatical features) (Carrasco-Ortíz et al., 2017). Since the target language in our experiment (Spanish) is typologically different from the participants’ native language (Japanese), and the specific morphosyntactic violations tested in our experiment have no Japanese analog, it is unclear how our participants might process the Spanish morphosyntactic violations. This question is further confounded by the generally low proficiency level of our participants, although the target grammar is well within their grasp. Taken together, we expect morphosyntactic violations to elicit an N400-like negativity in our L2 participants, but considering the linguistic distance between Spanish and Japanese, we do not rule out the possibility of some other neural strategy for morphosyntactic processing.

### 1.4 Expected Results: Orthographic/Phonological Violations

Within the context of this experiment, orthographic violations and phonological violations more specifically refer to misspellings and mispronunciations, respectively.

#### L1 Experiment

When considering what ERP component(s) to expect in the orthographic violation condition, many factors weigh in, including cloze probability, whether the misspelling is pseudohomophonic or not, whether or not the misspelling is an orthographic neighbor, the stress of the violated syllable, etc. (Kriukova & Mani, 2016; Laszlo & Federmeier, 2009; Newman & Connolly, 2004). In most cases, an N400 is elicited, and the aforementioned characteristics of the violation affect only its amplitude or latency, without resulting in an entirely different ERP component. Since our experiment violates the second word, which is preceded only by a subject pronoun, cloze probability is not a confounding variable. Instead, the orthographic/phonological violations in our experiment closely match those of Kryuchkova & Mani (2016), but in Spanish and without controlling for stress. Therefore, we similarly expect to elicit an N400 and/or P600 for our orthographically violated condition.

Although ERP data on the processing of mispronunciations is relatively scarce, previous research indicates that an N400 is to be expected for the phonologically violated (i.e. mispronounced) condition (Friedrich, Eulitz, & Lahiri, 2006). Since we did not control for the direction of the violation (e.g. coronal to non-coronal, or vice-versa), it is possible that some mispronunciations either went completely unnoticed or elicited smaller effects due to the underspecification of certain phonological features (Lahiri & Reetz, 2002). However, only trials with correctly identified violations were included in the ERP analysis, thereby minimizing such effects.

#### L2 Experiment

Although some studies are tangentially related (e.g., Kasparian & Steinhauer, 2016; van Rees Vellinga, Hanulíková, Weber, & Zwitserlood, 2010), we could not find any previous research directly investigating L2 processing of misspellings or mispronunciations. On the other hand, one study conducted on high-proficiency learners of Mandarin Chinese showed that vowel substitution resulting in nonwords elicited an N400, while tone substitutions resulting in nonwords did not, with behavioral accuracy favoring the former task over the latter (Pelzl, Lau, Guo, & Dekeyser, 2021). In light of these results, it seems fair to expect that mispronunciations resulting in nonwords will elicit an N400-like response in high-proficiency L2 learners who accurately recognize the violation as a nonword, as opposed to a word they simply have not yet learned. However, it seems likely that lower-proficiency learners would be unable to distinguish when a misspelling or mispronunciation is a nonword, given that all violations in our experiment resulted in phonotactically legal words in Spanish. In light of this, we expect L2 learner performance on this task to be poor and the resulting ERPs to lack any defining N400-like negativities.

## 2 Methods

### 2.1 Participants

#### L1 Experiment

Eighteen native Spanish speakers (mean age 28.0 ± 4.3 years, range 20.8–35.8; 15 males) participated in the study. All participants were right-handed (mean LQ 86.7 ± 20.3, range 35– 100) according to the Edinburgh Handedness Inventory (Oldfield, 1971) and had no history of neurological disorders. As the experiment was conducted at a Japanese university, all participants were living in Japan, came from different Spanish-speaking countries, and had varying degrees of proficiency in English and Japanese.

Participants were paid 1,000 JPY per hour of their participation, plus a 1,000 JPY travel stipend where applicable. Since the experiment consisted of two three-hour sessions, participants were generally paid 8,000 JPY in total.

#### L2 Experiment

Twenty-four native Japanese-speaking L2 Spanish learners (mean age 20.9 ± 1.7 years, range 18.9–26.7; 10 males) were recruited to participate in the L2 experiment.

Recruits were subjected to a screening session. During the screening session, participants took a proficiency test compiled from the written components of levels A1, A2, B1, and B2 of the *Diplomas de Español como Lengua Extranjera* (DELE, Diplomas of Spanish as a Foreign Language) sample tests, which were freely available through the DELE website at the time. These DELE levels are designed to roughly correspond to the Common European Framework of Reference for Languages (CEFR), which ascribes a proficiency level to L2 speakers from one of the following (ranked lowest to highest): A1, A2, B1, B2, C1, C2. This test was provided to assess overall Spanish proficiency. In addition to the DELE-based proficiency test, participants took a shortened version of both the visual and auditory experiments described in Section 2.2 in order to ensure that they were capable of performing the experimental tasks at a sufficient level. In order to pass the screening test and proceed to the experiment, participants were required to score better than 20% on the proficiency test, which roughly corresponded to an A1 level, and score at least 65% on the grammar judgment task.

Of the 24 recruits, 18 participants (mean age 20.46 ± 1.1 years, range 18.8–23.2; 9 males) passed the screening and completed the experiment in both modalities. Among those who passed, the average score on the proficiency test was 0.433 ± 0.145, and average score on the screening grammar judgment task was 0.811 ± 0.081. All participants were native Japanese speakers, right-handed (mean LQ = 90.3 ± 11.4, range 70–100) according to the Edinburgh Handedness Inventory (Oldfield, 1971), and had no history of neurological disorders.

The 18 L2 participants were divided into high and low proficiency groups based on their DELE-based proficiency screening test scores. The median performance on the DELE exam was used as the inclusive cutoff between the two proficiency groups, resulting in 10 high proficiency participants (mean DELE score = 0.529 ± 0.113) and 8 low proficiency (mean DELE score = 0.313 ± 0.123) participants.

### 2.2 Stimuli & Tasks

Since this study investigates both native and non-native language processing, stimuli was made to be accessible to both L1 and L2 speakers of Spanish. To that end, the first priority in constructing stimuli was the use of vocabulary words that were covered in a first-semester Spanish class at Kyushu University, where the participants attended university.

Stimuli consisted of 240 Spanish sentences, all of which followed canonical Spanish structure: Subject-Verb-Object. The target 120 Spanish sentences were comprised of three conditions: control, morphosyntactic violation, orthographic/phonological violation (see Table 1). The remaining 120 sentences were unviolated filler sentences. Tenses covered in the first semester of Spanish (present indicative, preterite, imperfect indicative, and future indicative) were included and their frequency was matched to the relative distribution of natural Spanish (Bull et al., 1947), in order to obscure experiment purposes with more natural stimuli. Only present-tense verbs were violated. Although Spanish is a pro-drop language, which allows subject pronouns to be omitted, the subject was always explicitly given. The 2nd-person plural (*vosotros*) form was omitted when constructing stimuli since L1 participants came from various countries and this form is not used universally among Spanish-speaking countries.

**Table 1.**
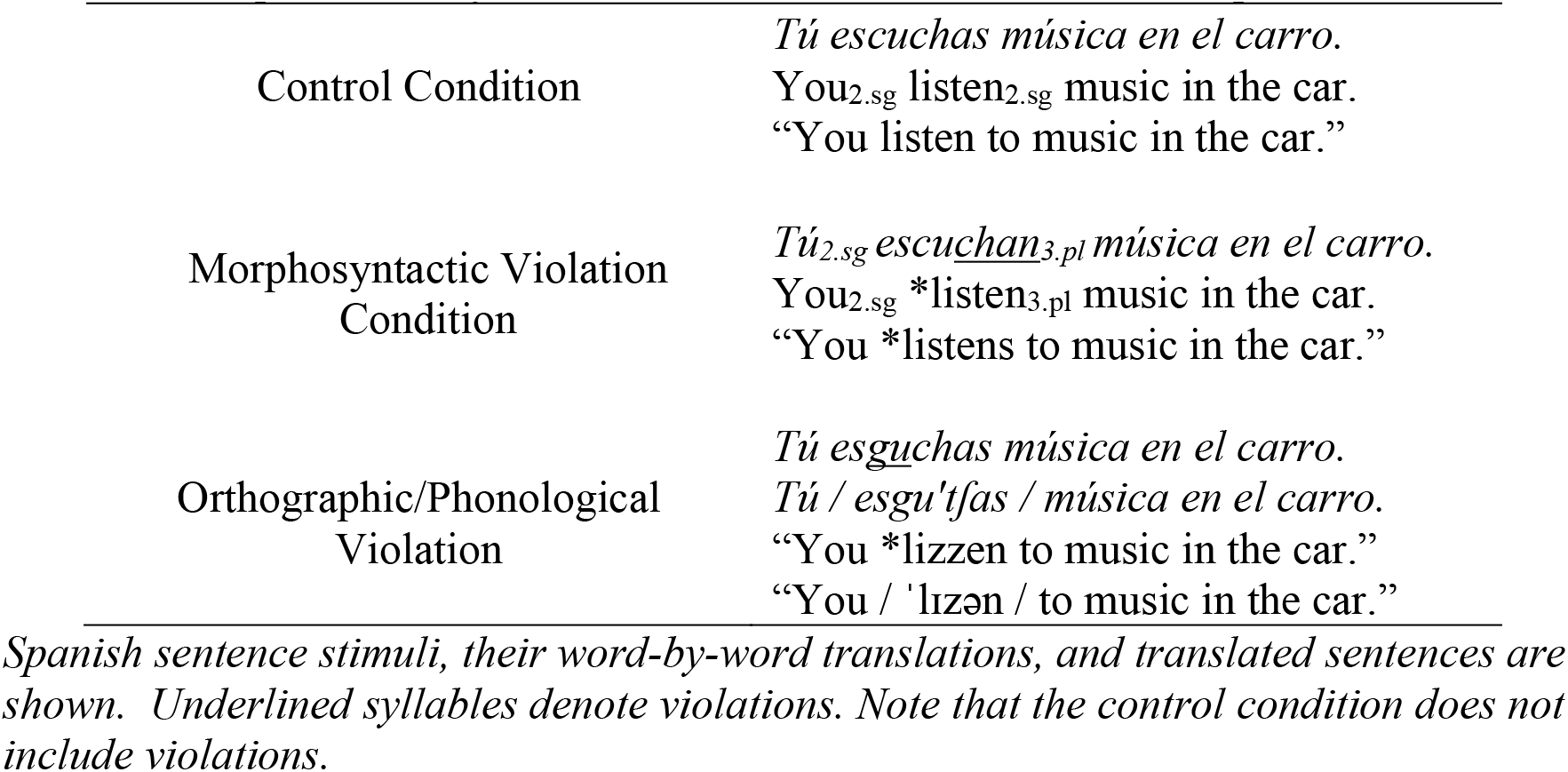
Example stimuli of each condition used in both L1 and L2 experiments.

In the morphosyntactic violation condition, the verb was conjugated to disagree with the immediately preceding explicit subject in person, number, or both. The pairing of disagreements between subject person/number and verb person/number was evenly distributed across all possible combinations. However, since Spanish allows for Unagreement (pairing a first-person plural verb with a third-person plural noun), and since this pairing is processed as an unviolated verb (Mancini, Molinaro, Rizzi, & Carreiras, 2011b), this particular pairing was omitted.

In the orthographic/phonological violation condition, the verb spelling/pronunciation was changed to a minimal pair pseudoword. The minimal pair was constructed in a manner similar to Kriukova and Mani (2016), by replacing one consonant with another consonant that differed by only one distinctive phonological feature (for example by adding or subtracting voicing [± voice]).

Sentences were randomized and paired with one of three tasks: comprehension task (e.g., “where do you listen to music?”), grammatical judgment task (“was the preceding sentence grammatically correct?”), and orthographic/phonological judgment task (“was the preceding sentence spelled/pronounced correctly?”). The comprehension task questions were multiple-choice with two options, while the other two tasks were yes/no questions. In the L1 experiment, tasks were presented in Spanish, while in the L2 experiment, tasks were presented in Japanese. Control condition stimuli and filler sentences were paired with all task types, while morphosyntactically violated stimuli and orthographically/phonologically violated stimuli were always paired with their respective judgment tasks. In doing so, we were able to make answers evenly split between the two options (i.e., A/B multiple choice for comprehension task and yes/no for judgment tasks).

Figure 2 illustrates how stimuli were presented. As shown, stimuli began with fixation crosses, followed by either a blank screen in the visual experiment or continued silence in the auditory experiment (i.e., the inter-stimulus interval, ISI). The subject was presented for a duration matching the auditory stimulus duration. A variable ISI followed to ensure that all sentences had a consistent verb onset of 1032 ms, which was decided by the longest-duration subject pronoun (*‘nosostros’,* ‘we’) plus a 100-ms ISI (see Barry, Fogarty, De Blasio, & Karamacoska, 2018 for reductive effects of variable ISI on preferential phase occurrence). After the critical word, a constant ISI of 100 ms was included between each word/phrase.

**Figure 2.**
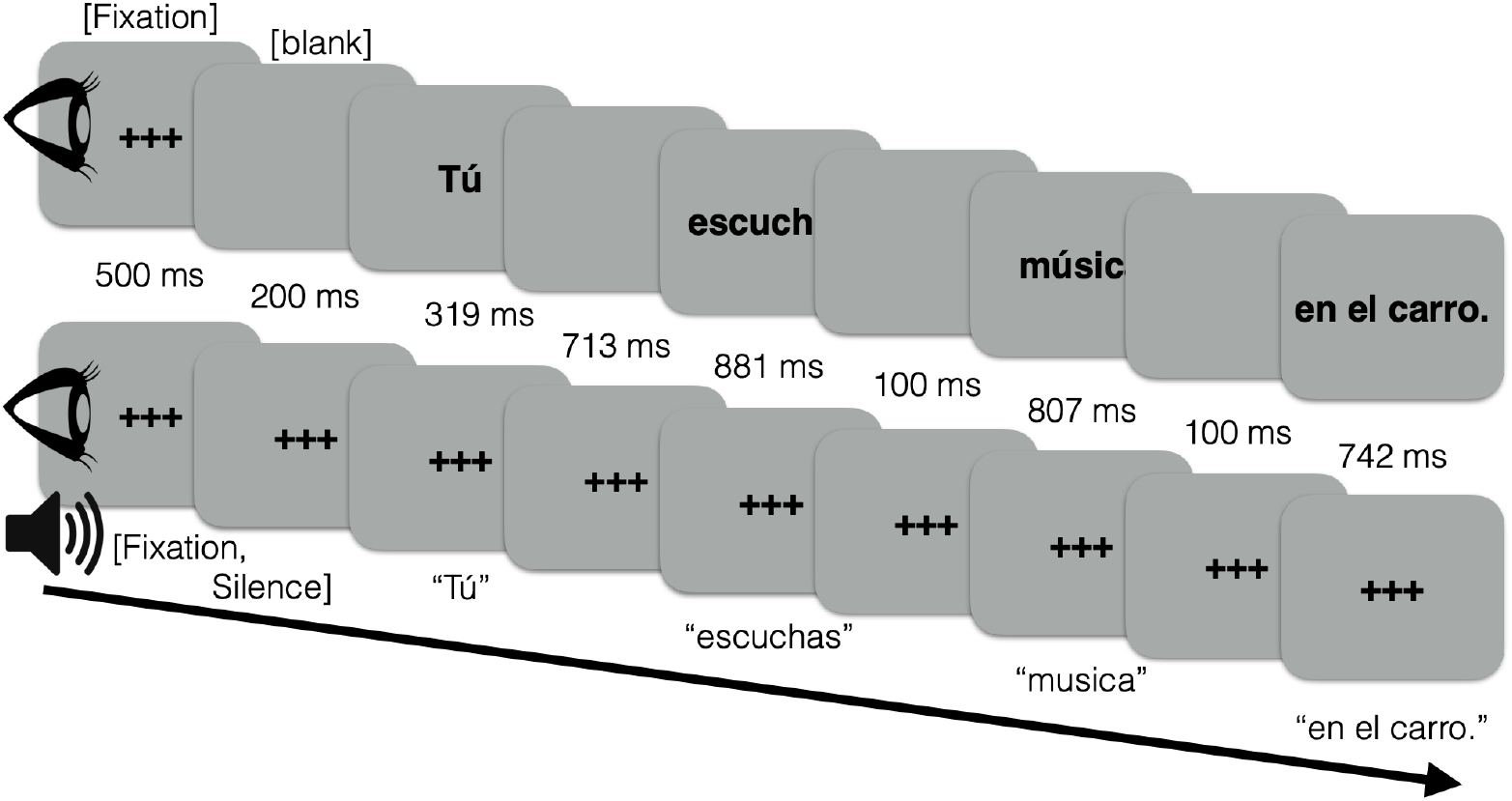
A schematic diagram illustrating stimulus presentation in the visual (top) and auditory (bottom) modalities. Subjects and verbs were presented individually, and thereafter, depending on content, sentences were presented by word or phrase. Durations of each Spanish word/phrase in visual stimuli were the same as the auditory stimuli durations.

Stimuli were presented and behavioral data was acquired using Presentation® software (Version 18.0, Neurobehavioral Systems, Inc., Berkeley, CA, www.neurobs.com). Statistical analysis of behavioral data was performed in Microsoft Excel, Python 3.7.0, and R 3.5.1.

### 2.3 EEG Setup

64 active Ag/AgCl electrodes were attached to the participant’s scalp for measuring electrophysiological activity in a montage based on an extended international 10-20 localization system.

### 2.4 ERP Analysis

EEG data processing was done in MATLAB with the MATLAB toolbox EEGLAB, and in Python with the open-source toolbox MNE-Python.

EEG data was recorded online with a single reference electrode on the right ear lobe. In order, the steps for computing the grand average ERPs included manual removal of bad channels, bad channel interpolation, applying a high-pass filter at 0.1 Hz, re-referencing to the whole-scalp average (i.e., common average reference), independent component analysis (ICA) and IC removal, epoching, baseline correction using a 200 ms time-window pre-stimulus onset, rejecting artifacts more than ±100 μV relative to the baseline, applying a low-pass filter at 30 Hz, and taking the weighted average of all ERPs.

Visual evoked ERP data was aligned to critical word onset (i.e., verbs), while auditory evoked ERP data was aligned to critical syllable onset. As mentioned in Section 1.2, auditory information is revealed across time, which is variable depending on the length of the word, the speaker’s speech rate, etc. This in turn results in a variable timing of exposure to the violation, which in our experiment ranged from 492 ms (*hablas*) to 1126 ms (*manejamos*) post word onset in the morphosyntactic violation condition and from 433 ms (*funa*) to 1226 ms (*disfrudamos*) post word onset in the phonological violation condition. It was often the case that exposure to the violation occurred more than 600 ms post word onset. Thus, using word onset would necessarily obscure the timing of the P600 and other ERP components observed. Ideally, word length is controlled (Hauk & Pulvermüller, 2004), but our experimental stimuli required prioritizing intelligibility among low-level L2 learners, substantially limiting the pool of verbs to be drawn from, and precluding the option of controlling word length. However, the use of critical syllable onset to align auditory evoked ERPs is not novel (see Baart & Samuel, 2015; Kooijman, Hagoort, & Cutler, 2009) and it provides a much more consistent delay between the stimulus onset and violation. Thus, each auditory stimulus sound file was measured for the precise timing of the violated syllable in each violation condition and its corresponding unviolated syllable in the control condition.

**Figure 3:**
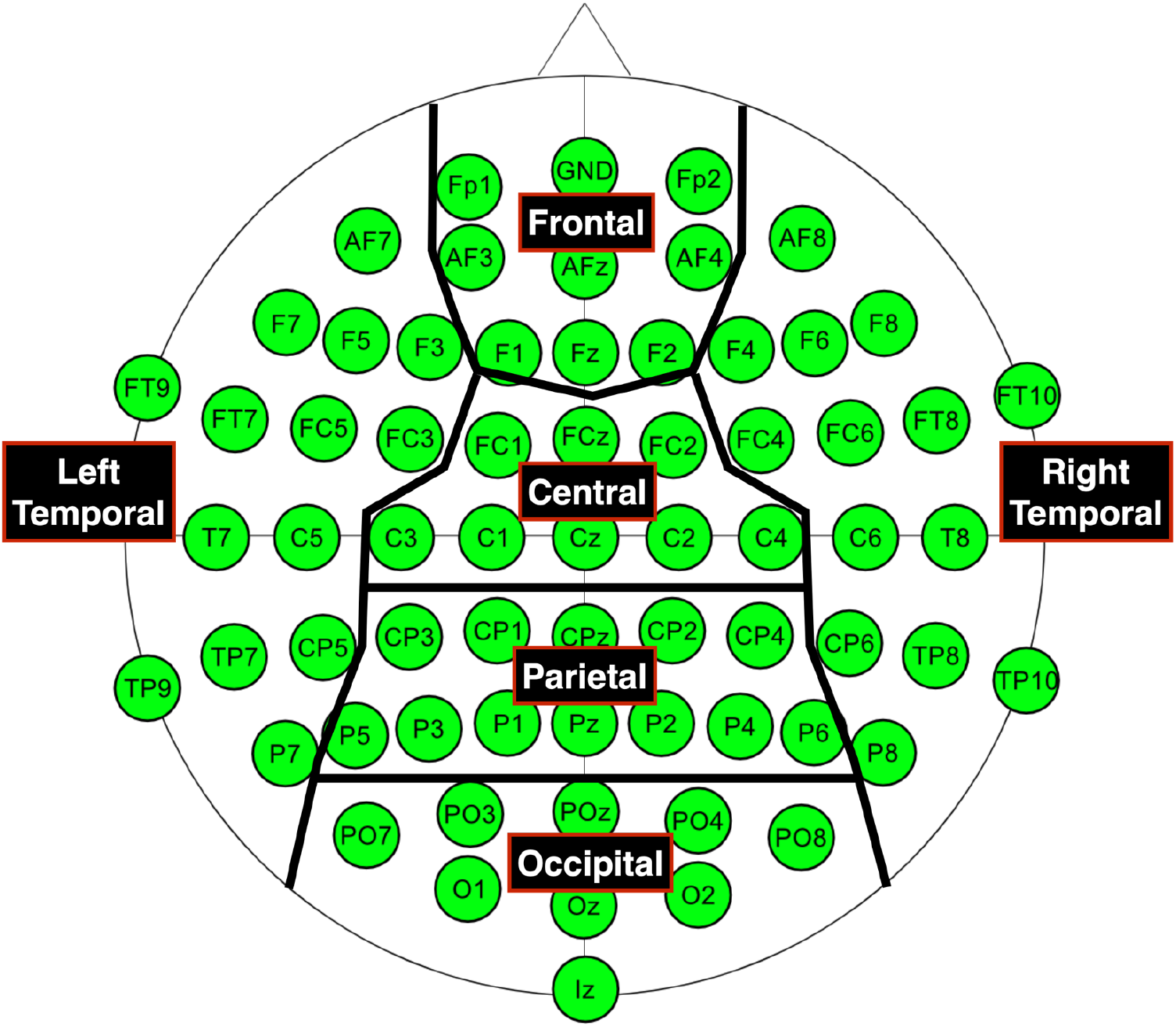
Six regions across the scalp used for statistical analysis.

For statistical analysis of ERP data, the scalp electrodes were divided into six regions, as shown in Figure 3. Statistical analysis of ERP data was performed in Microsoft Excel, Python 3.7.0, and R 3.5.1.

By visually inspecting topographical maps of voltage distribution across the whole epoch, we compiled lists of ROIs to test for significance using LME models. To test for a modality effect, we looked for significant interactions between modality and condition, since this would correspond to a difference in processing between conditions that is contingent upon the modality of the stimulus.

To mitigate the family-wise error rate (FWER) that arises with multiple comparison testing, we implemented Holm’s Sequential Bonferroni Procedure to calculate the adjusted p values.

## 3 Results

### 3.1 L1 Experiment

#### 3.1.1 L1 Behavioral Results

Task accuracies and response times (RTs) are shown in Figures 4 and 5, respectively. Two-sided paired t-tests assuming equal variance were run on accuracies and RTs to check for a significant effect of modality.

**Figure 4.**
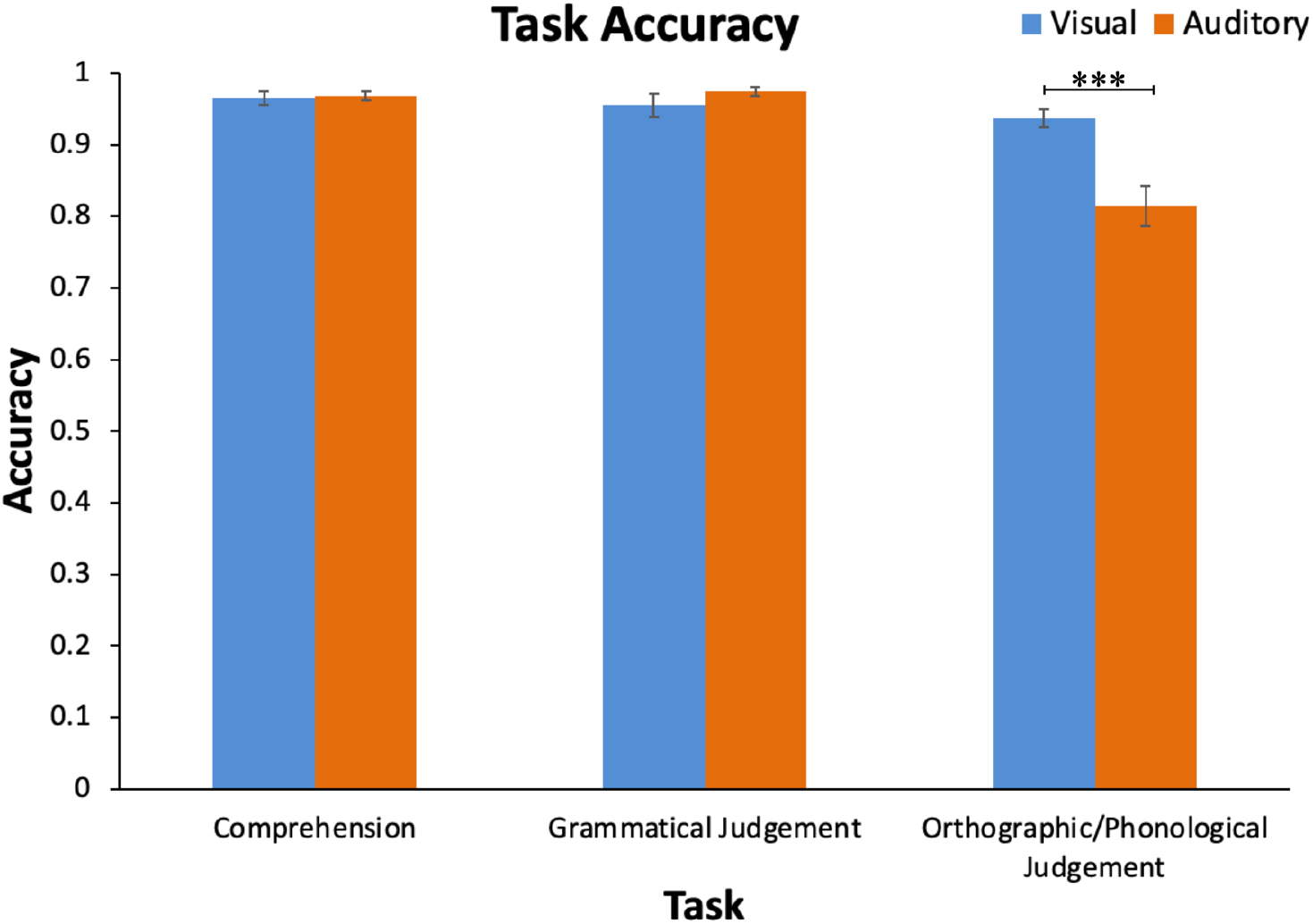
L1 accuracies in each modality (visual: blue, auditory: orange). Error bars represent standard error. ***p<0.001

**Figure 5.**
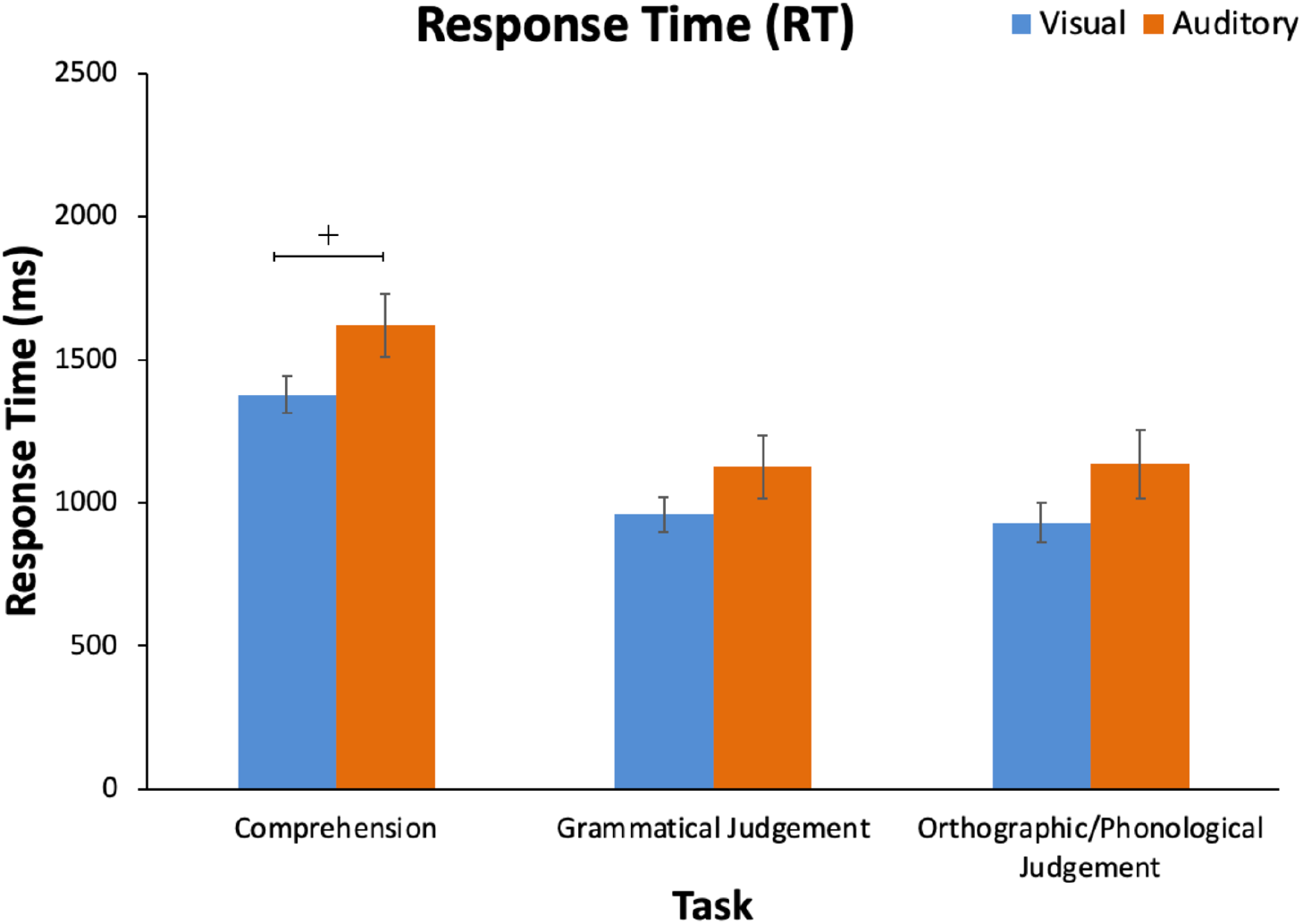
L1 response times in each modality (visual: blue, auditory: orange). Error bars represent standard error.

Participants were significantly less accurate in the phonological judgment task in the auditory modality (M = 0.814, SD = 0.121) than the orthographic judgment task in the visual modality (M = 0.938, SD = 0.054, *t*(34) = 3.969, *p* < 0.001). There was no significant effect of modality in the comprehension or morphosyntactic judgment tasks (*t*(34) = −0.236, *p* = 0.815 and *t*(34) = −1.079, *p* = 0.288, respectively), with all accuracies approaching 1 (visual comprehension M = 0.965, SD = 0.043; auditory comprehension M = 0.968, SD = 0.025; visual morphosyntactic judgment M = 0.956, SD = 0.071; auditory morphosyntactic judgment M = 0.975, SD = 0.023).

No significant effect of modality was observed in any task’s response time, although the comprehension task was marginally significant (*t*(34) = −1.885, *p* = 0.068), with the visual modality response time (M = 1377.9 ms, SD = 272.3 ms) being faster than the auditory modality response time (M = 1619.5 ms, SD = 470.6 ms). Although, visual responses were all faster than auditory response times (visual morphosyntactic M = 959.3 ms, SD = 262.3 ms; auditory morphosyntactic M = 1125.3, SD = 466.2 ms; visual orthographic violation M = 930.1 ms, SD = 292.2 ms; auditory phonological violation M = 1136.6 ms, SD = 507.4 ms), none of these difference rose to the level of significance (morphosyntactic violation *t*(34) = −1.317, *p* = 0.197; orthographic/phonological violation *t*(34) = −1.496, *p* = 0.144).

#### 3.1.2 L1 Electrophysiological Results

Single-participant data with fewer than 13 trials in any given condition remaining after artefact rejection were excluded from all conditions. This resulted in the exclusion of two visual datasets and four auditory datasets. Thus, ERP data was analyzed from 16 visual participants and 14 auditory participants.

ERP data was analyzed using difference waveforms and difference topography, which were computed by subtracting the control condition from either the morphosyntactic violation condition or the orthographic/phonological violation condition. Since we are primarily investigating a neurophysiological modality effect in language processing, using this difference condition allows us to remove sensory perceptual effects and other non-linguistic processing that may contribute to modality-specific EEG recordings, but are not of interest.

#### 3.1.3 L1 Electrophysiological Results – Morphosyntactic Violations

As described in Section 1, morphosyntactic violations robustly elicit a P600 in both modalities. Due to their ubiquity in neurolinguistics literature, we start with a discussion the N400 and P600 elicited by morphosyntactic violations, shown in Figure 6.

**Figure 6:**
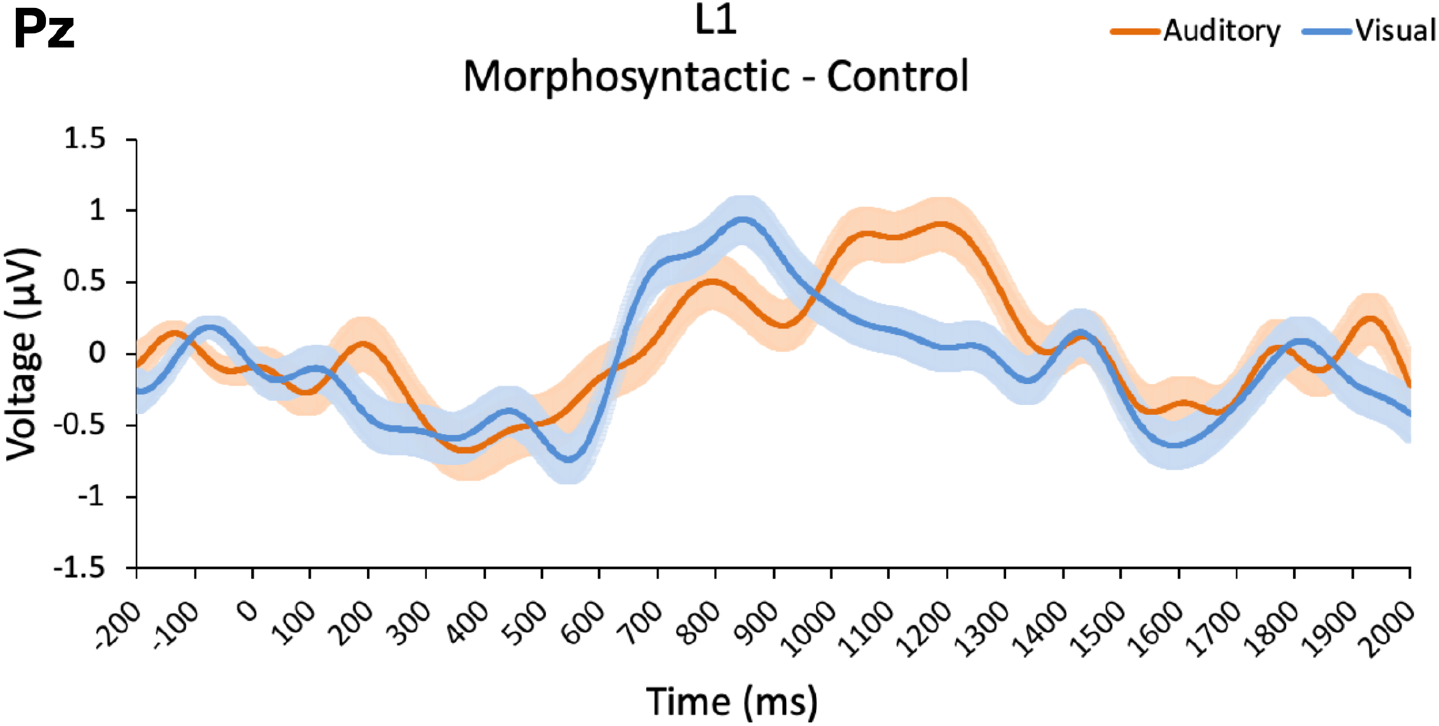
Morphosyntactic violation difference waveform (morphosyntactic violation minus control) showing the P600 in both modalities over Pz. For display purposes, the waveform is plotted with a low-pass filter at 6 Hz. Positivity is plotted upward.

As can be seen in Figure 6, the two modalities overlap in most places. A small N400 is exhibited in both modalities, followed by a strong P600. In the auditory modality the positive deflection is more gradual and peaks after the visual modality (auditory peak at 1188 ms; visual peak at 847 ms). It is also noteworthy that the visual evoked P600 diminishes substantially even before the auditory evoked P600 has peaked. Despite the differences in peak latency, both the auditory evoked and visual evoked P600s go positive (auditory 671 ms; visual 633 ms) and negative (auditory 1474 ms; visual 1468 ms) with similar timing. Additionally, the two modalities have comparable peak amplitudes, with the auditory evoked P600 peaking at 0.907 μV and the visual evoked P600 peaking at 0.941 μV. Relevant statistical analyses are provided in Table 2, along with analyses of other components.

**Table 2:** P values of ROIs and their corresponding time windows in the L1 morphosyntactic violation difference condition using the LME model Amplitude ~ Modality * Condition + (1|Subject).

| Region | Time (ms) | Modality | Condition | Modality:Condition | Modality:Condition (adj.) |
| --- | --- | --- | --- | --- | --- |
| Occipital | 600–700 | 0.2022 | 0.0932<sup>+</sup> | 0.0998<sup>+</sup> | 0.0998<sup>+</sup> |
| Frontal | 600–700 | 0.0029** | 0.11256 | 0.08671<sup>+</sup> | 0.17342 |
| L. Temporal | 1100–1200 | 6.51E-05*** | 0.6166 | 0.007** | 0.035* |
| R. Temporal | 1100–1300 | 0.00126** | 0.57851 | 0.07163<sup>+</sup> | 0.21489 |
| Parietal | 1100–1300 | 9.93E-07*** | 0.9952 | 0.0185* | 0.074<sup>+</sup> |

Adjusted *p* values were calculated by the Holm-Bonferroni method. <sup>+</sup>*p* < 0.10, \**p* < 0.05, \*\**p* < 0.01, \*\*\**p* < 0.001

One might consider that using critical syllable onset to align auditory ERPs is an imperfect solution to account for the information delay in the auditory modality (while the stimulus unfolds in time), and therefore causes a latency shift in the auditory evoked P600. For this reason, we also include a peak-aligned ERP waveform, shown in Figure 7, where the auditory evoked ERP was shifted to align the auditory P600 peak with the visual P600 peak (as calculated by the average of all parietal electrodes), resulting in shifting the auditory evoked ERP 344 ms earlier. The waveform was additionally recorrected for its new baseline, slightly shifting the whole waveform up (by 0.222 μV). This peak-alignment method actually leads to a greater difference between the two modalities, namely that the auditory evoked ERP starts going positive much earlier (at 327 ms) than the visual evoked ERP.

**Figure 7:**
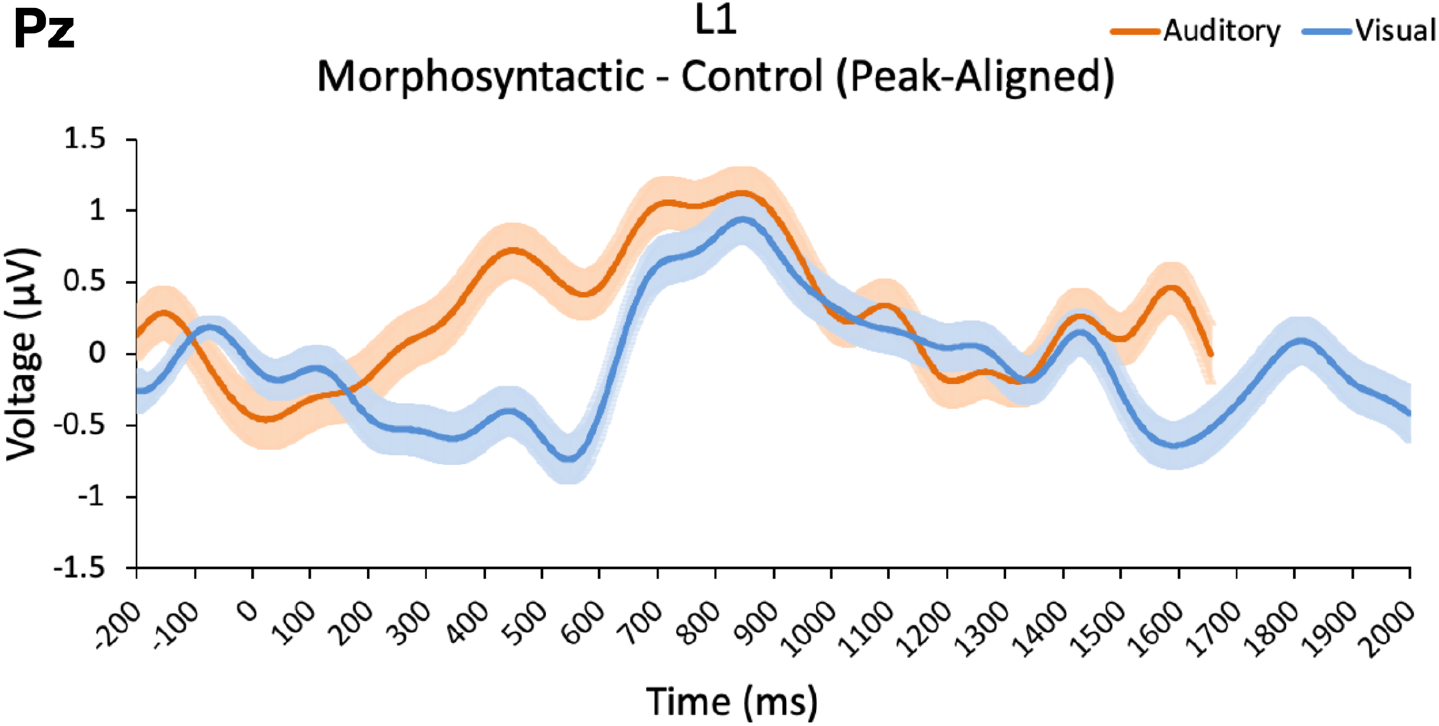
Peak-aligned difference waveform (morphosyntactic violation minus control) at Pz. Auditory waveform was epoched up to 2000 ms post critical syllable onset but was shifted to align the auditory P600 peak with the visual P600 peak, using the average of all parietal electrodes. After realignment, the auditory waveform only had data up to 1656 ms (2000 ms - 344 ms) and was recorrected with its new 200 ms baseline. For display purposes, the waveform is plotted with a low-pass filter at 6 Hz. Positivity is plotted upward.

As we see in Figure 7, regardless of the temporal alignment chosen, the P600 ERP component exhibits some modality-specific characteristics. Considering the two alignments holistically, we consider the use of the critical syllable onset to be the prudent choice, as it results in greater overlap between the two ERP components, especially the overlapping of the N400 component, than does the peak-aligned method. Ultimately, the temporal discrepancy between visual and auditory stimuli may have no perfect solution, but the data shown in Figure 6 does suggest that critical syllable onset is a viable approach for comparing the two modalities.

In this study, we are not limiting our investigation to the P600. Instead, we take an exploratory approach using topographical maps of voltage distribution to look for anomalous data throughout the whole ERP epoch (2000 ms). We first take a qualitative look at the morphosyntactic violation difference condition in each modality separately, as shown in Figure 8.

**Figure 8:**
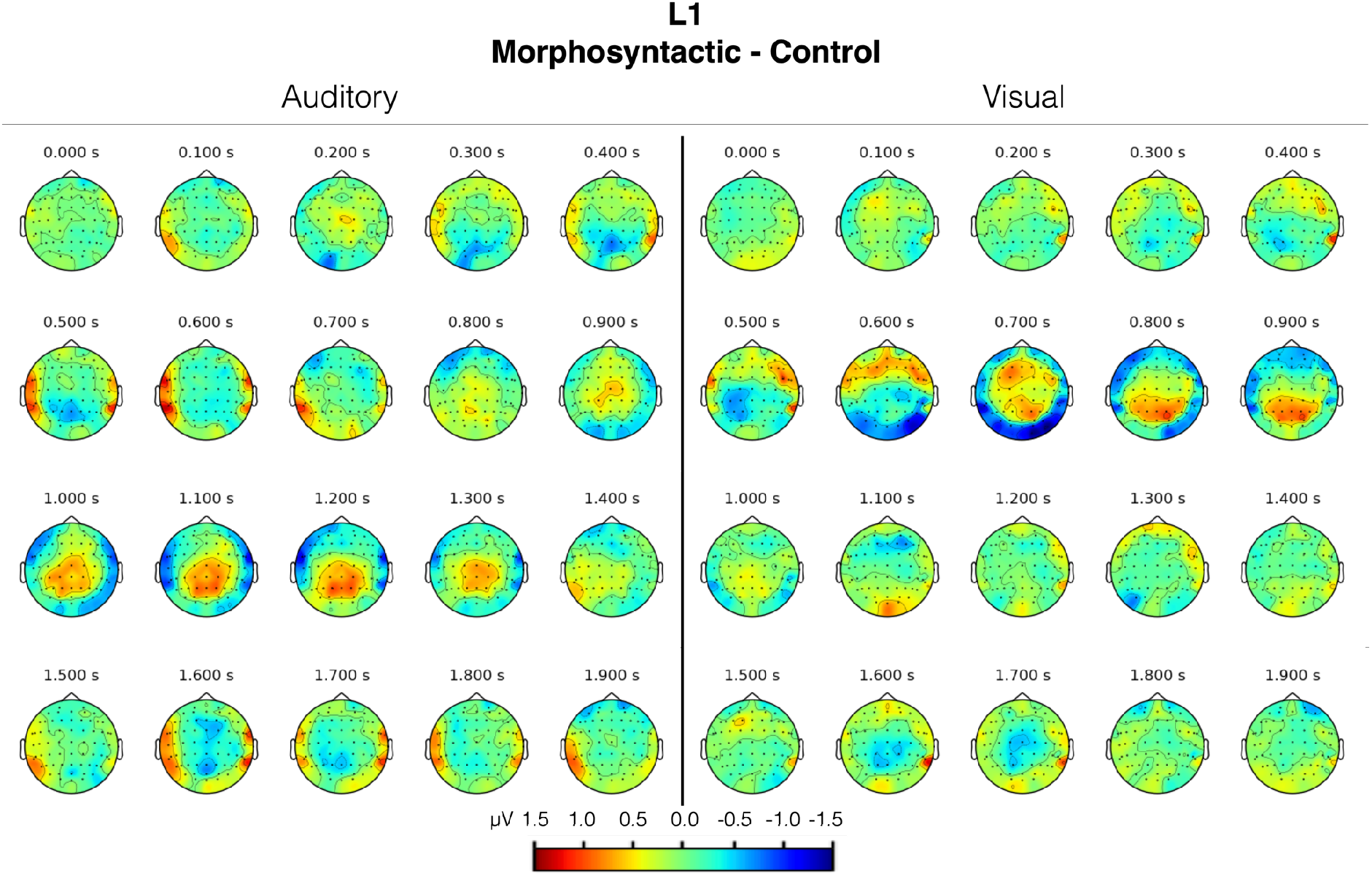
Topographical maps of voltage distribution for the morphosyntactic violation difference condition in the visual and auditory modalities. Each 100 ms interval is shown as a moving average with a 100 ms time window.

Figure 8 reinforces the data shown in Figure 6, displaying biphasic parietal negativities and positivities in both modalities. Again, we see the auditory positivity being more gradual and peaking later than the visual positivity. Other activity in the temporal and occipital regions is also apparent. It is, however, easier to investigate the differences in modality by subtracting one from the other. In Figure 9, we have produced a “modality difference” topographical map, computed by subtracting the visual morphosyntactic violation difference condition from the auditory morphosyntactic violation difference condition. This “morphosyntactic violation modality difference” topographical map shows the (native) neurophysiological modality effect that is the subject of this study. Thus, we use this plot to select our regions of interest (ROIs).

**Figure 9:**
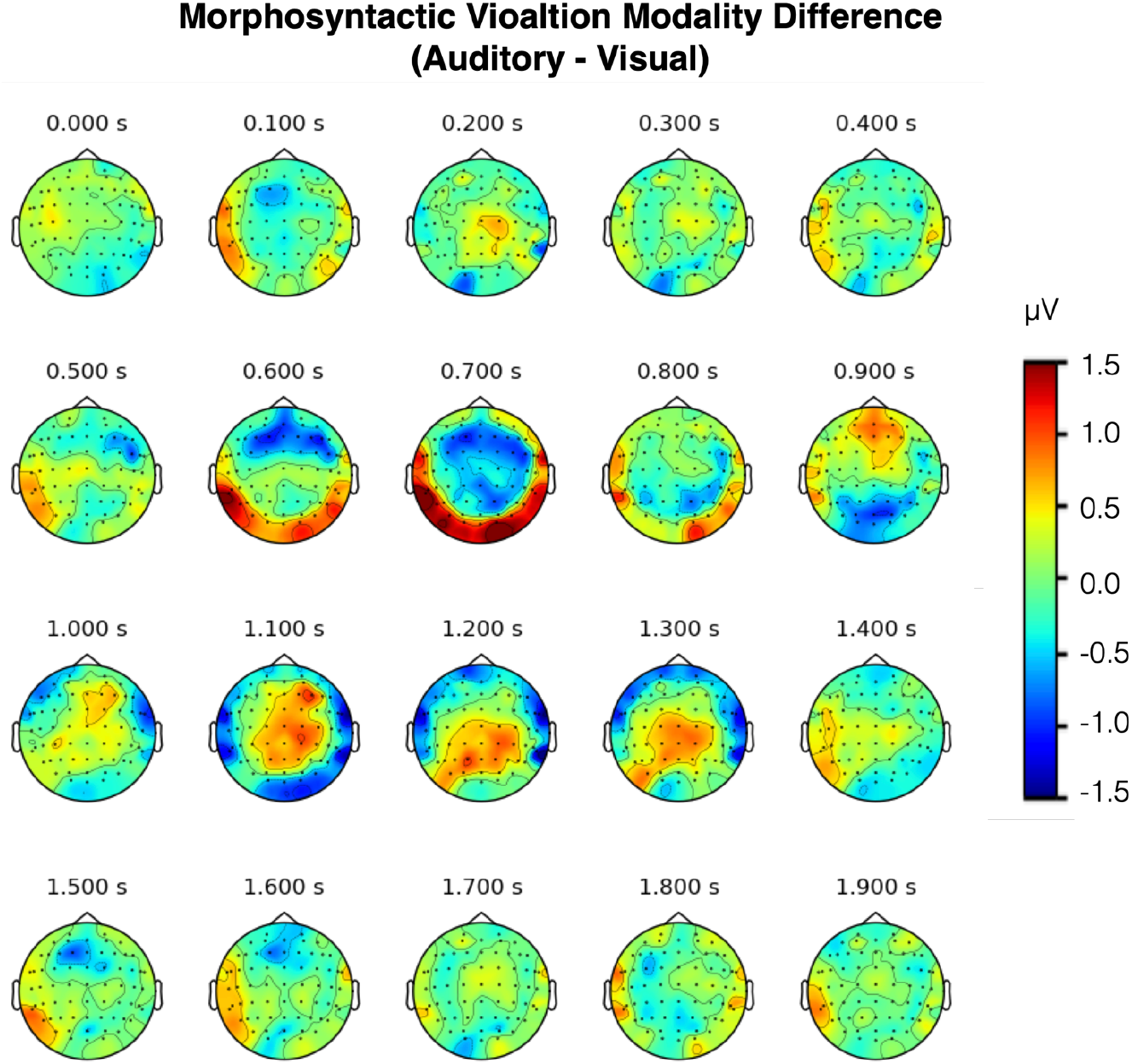
Topographical map of voltage distribution of the modality effect in L1 morphosyntactic violation processing across time. Each 100 ms interval is shown as a moving average with a 100 ms time window. Voltage was computed using the morphosyntactic violation difference condition of each modality and subtracting the visual from the auditory.

By visual inspection of Figure 9, we compiled a list of 5 ROIs, shown in Table 2, to test for a significant interaction between modality and condition, to indicate a neurophysiological modality effect in native morphosyntactic processing. Significance was tested using the LME model Amplitude ~ Modality * Condition + (1|Subject).

Table 2 shows that one ROI rose to the level of significant interaction between modality and condition, while two ROIs were shown to be approaching significance. The one significant ROI can be described as a left temporal negativity at 1100–1200 ms present only in the auditory modality (Figure 10E). It is worth noting that the left temporal region includes the primary auditory cortex as well Wernicke’s area (superior temporal gyrus), a language center of the brain implicated in language comprehension. The two marginally significant ROIs correspond to a visual negativity at 600–700 ms (Figure 10D) over occipital electrodes (where the primary visual cortex is located) and an auditory positivity over parietal electrodes 1100–1300 ms (Figure 10C), which corresponds to the later auditory P600 peak shown previously in Figure 6.

**Figure 10:**
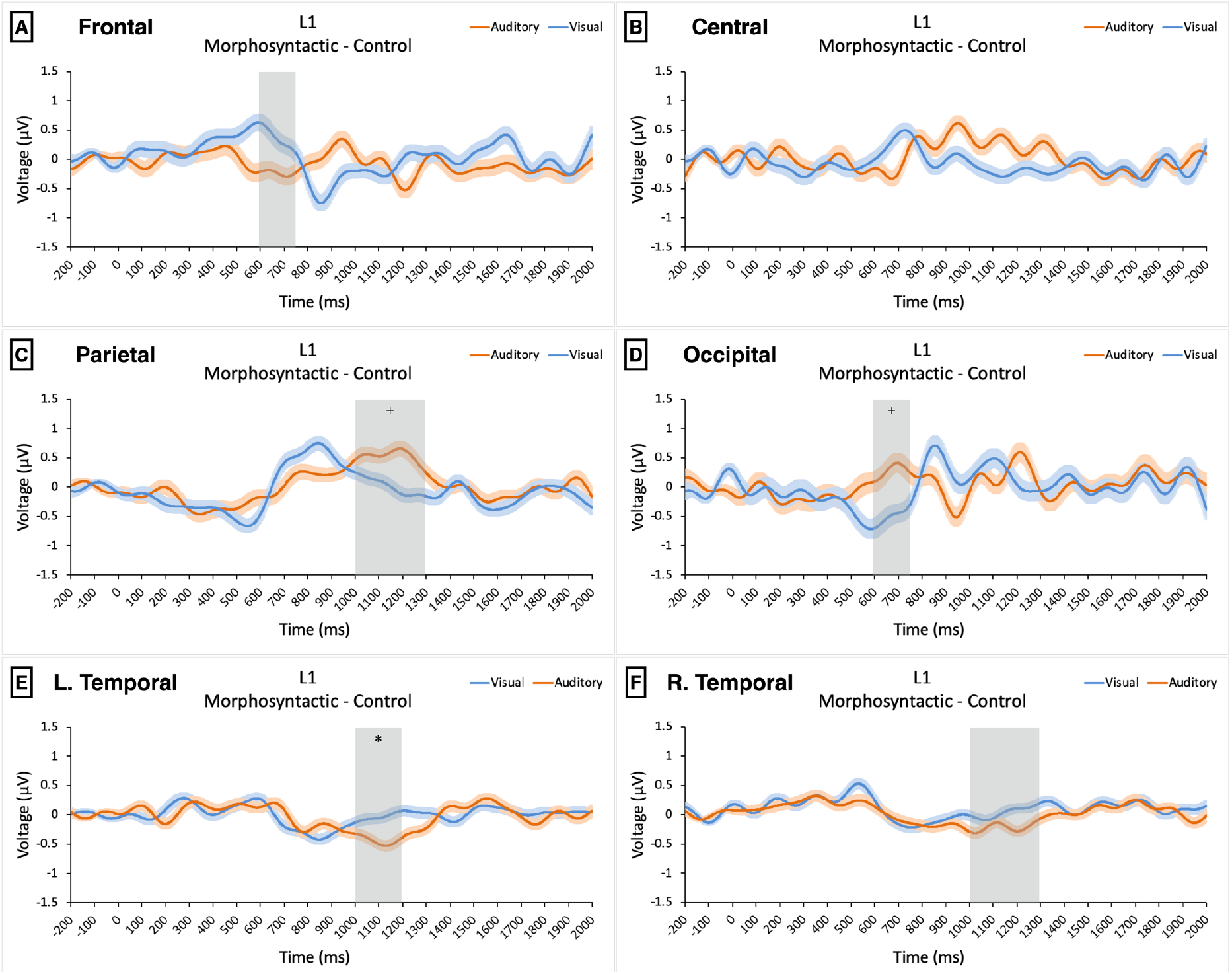
Morphosyntactic violation difference condition ERP waveforms for each region in L1 speakers. Each waveform is the average of all electrodes in the region shown. For display purposes, waveforms are plotted with a low-pass filter at 6 Hz. Positivity is plotted upward. Orange lines represent auditory evoked ERPs, while blue lines represent visual evoked ERPs. Standard error bars are included for each modality. The time windows of tested ROIs are highlighted in gray. ^+^p <0.10, *p < 0.05, all Holm-Bonferroni corrected.

The significant effects shown in Table 2 and Figure 10 represent evidence for a modality effect, demonstrating that morphosyntactic violation processing is comprised of both modality-dependent and modality-independent processing. The broader implications of this effect are discussed further in Section 4.1.

#### 3.1.4 L1 Electrophysiological Results – Orthographic/Phonological Violations

The analysis of orthographic/phonological violation-elicited ERPs was carried out in the same way as morphosyntactic violation-elicited ERPs in 3.1.3. We start with a qualitative examination of the whole-epoch topographical maps of voltage distribution in both modalities shown in Figure 11.

**Figure 11:**
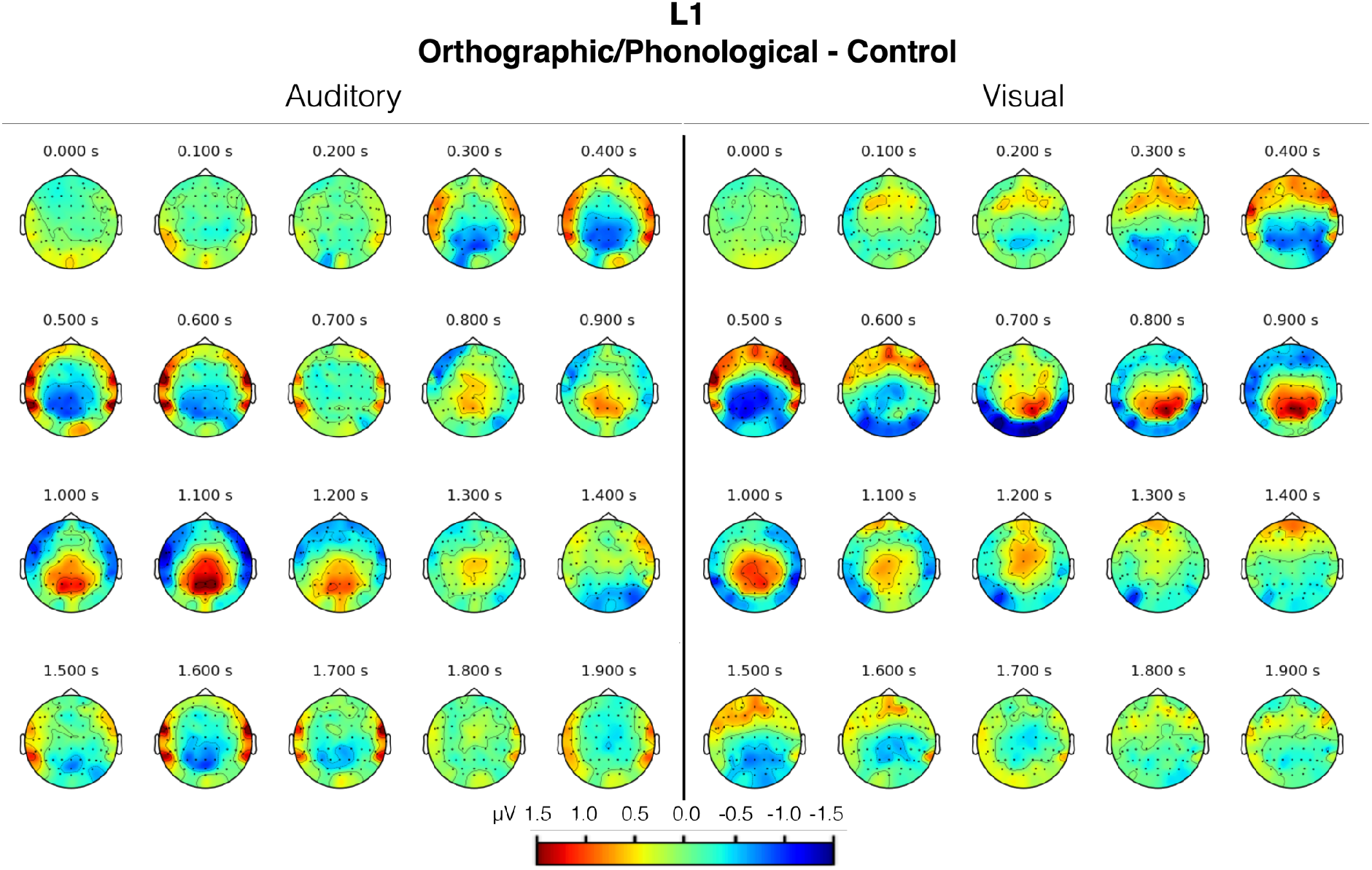
Topographical maps of voltage distribution for the orthographic/phonological violation difference condition in the visual and auditory modalities. Each 100 ms interval is shown as a moving average with a 100 ms time window.

A visual inspection of Figure 11 reveals a very similar neurophysiological response to morphosyntactic violation-elicited ERPs, namely a biphasic N400-P600 complex. Interestingly, just as was the case in the morphosyntactic violation condition, the N400 latency and magnitude are nearly identical across modalities, whereas the P600 is again more gradual in slope and later to peak in the auditory than in the visual modality.

One notable difference between the orthographic/phonological violation condition and the morphosyntactic violation condition ERPs is the magnitude of the N400, where the former elicits much greater negativities (in both modalities) than the latter. This may be due to increased difficulty of retrieval of misspelled and mispronounced words, whereas incorrectly inflected words should not elicit as much difficulty in the retrieval process.

To more precisely analyze modality-specific differences in the orthographic/phonological difference condition, we look at the orthographic/phonological violation modality difference topographical maps of voltage distribution shown in Figure 12.

**Figure 12:**
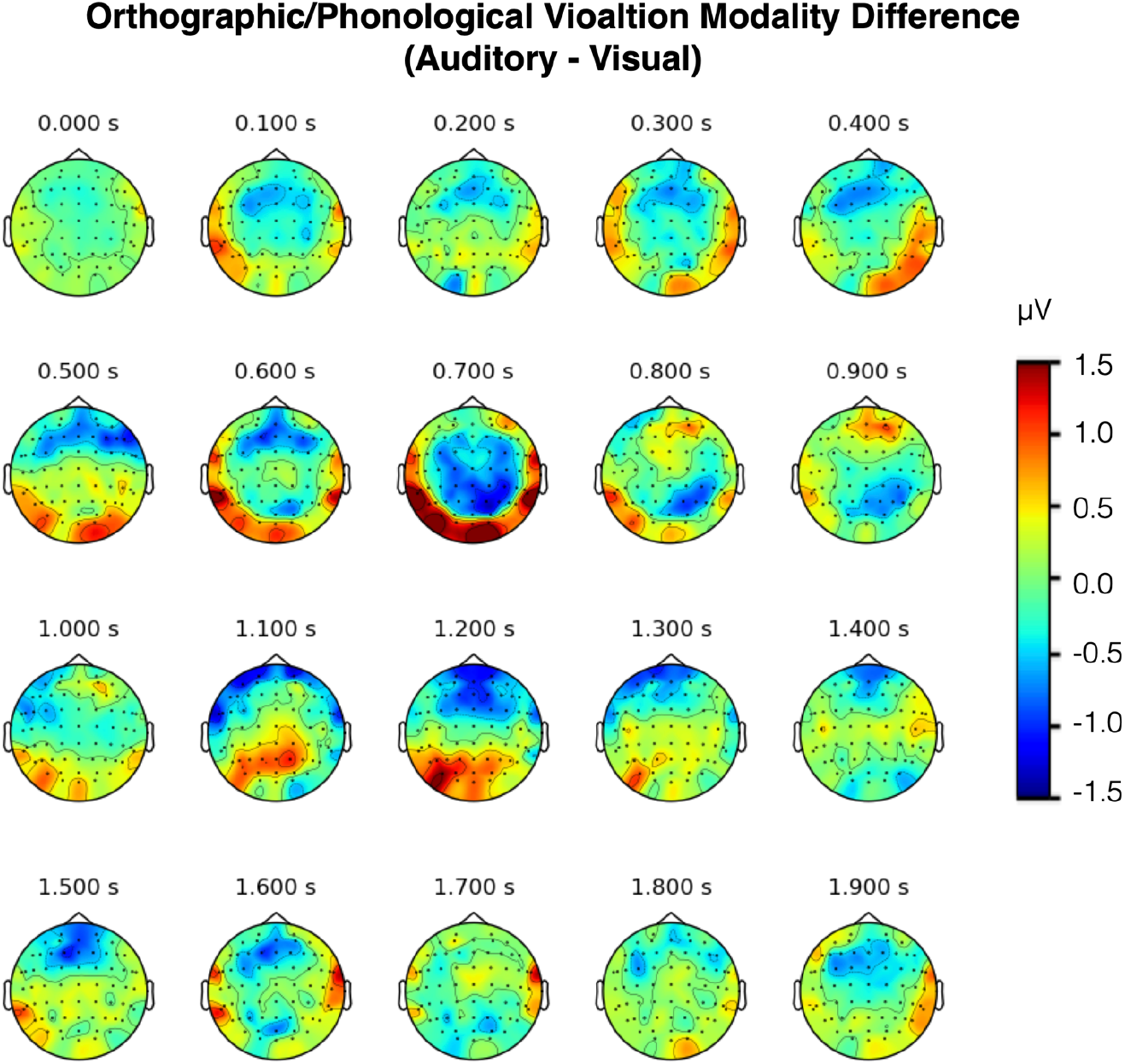
Topographical map of voltage distribution of the modality effect in L1 orthographic/phonological violation processing across time. Each 100 ms interval is shown as a moving average with a 100 ms time window. Voltage was computed using the orthographic/phonological violation difference condition of each modality and subtracting the visual from the auditory.

As was done in the morphosyntactic violation difference condition, by visual inspection of Figure 12, we compiled a list of 6 ROIs, shown in Table 3. We tested each ROI for a significance interaction of modality and condition using the same LME model (Amplitude ~ Modality * Condition + (1|Subject)) as before.

**Table 3:**
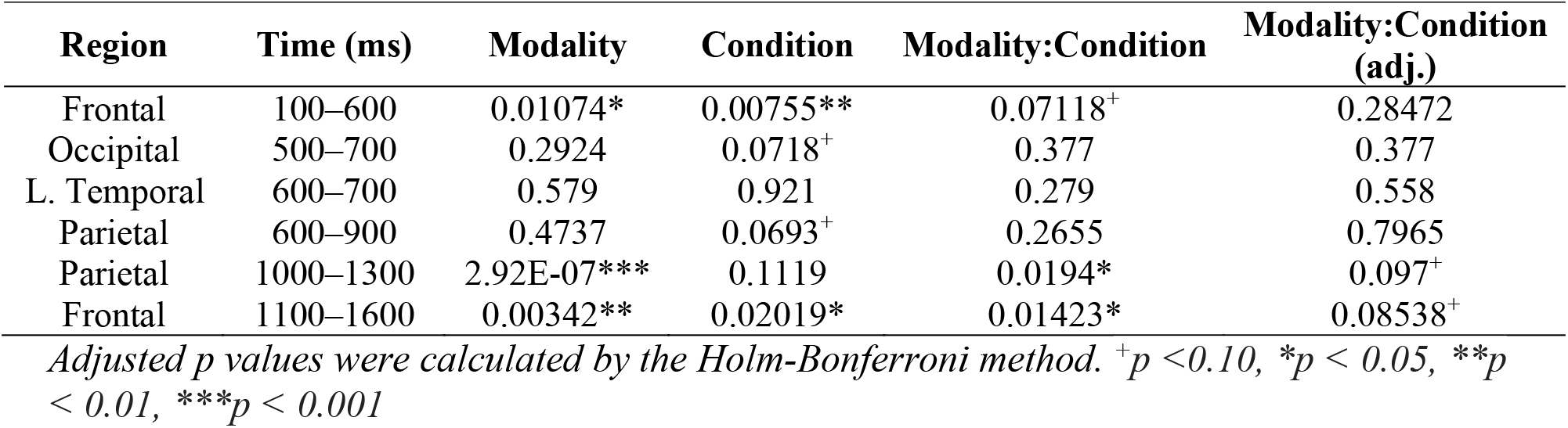
P values of ROIs and their corresponding time windows in the L1 orthographic/phonological violation difference condition using the LME model Amplitude ~ Modality * Condition + (1|Subject).

| Region | Time (ms) | Modality | Condition | Modality:Condition | Modality:Condition (adj.) |
| --- | --- | --- | --- | --- | --- |
| Frontal | 100–600 | 0.01074* | 0.00755** | 0.07118<sup>+</sup> | 0.28472 |
| Occipital | 500–700 | 0.2924 | 0.0718<sup>+</sup> | 0.377 | 0.377 |
| L. Temporal | 600–700 | 0.579 | 0.921 | 0.279 | 0.558 |
| Parietal | 600–900 | 0.4737 | 0.0693<sup>+</sup> | 0.2655 | 0.7965 |
| Parietal | 1000–1300 | 2.92E-07*** | 0.1119 | 0.0194* | 0.097<sup>+</sup> |
| Frontal | 1100–1600 | 0.00342** | 0.02019* | 0.01423* | 0.08538<sup>+</sup> |

Adjusted *p* values were calculated by the Holm-Bonferroni method. <sup>+</sup>*p* < 0.10, \**p* < 0.05, \*\**p* < 0.01, \*\*\**p* < 0.001

Table 3 shows that two ROIs were marginally significant (parietal 1100–1200 ms, Figure 13C; and frontal 1100–1600 ms, Figure 13A) and one ROI approaching significance (frontal 100–600 ms, Figure 13A). One of these ROIs corresponds to modality-specific P600 differences similar to those seen in the morphosyntactic violation difference condition, i.e., a more gradual and later-peaking positivity in the auditory modality (Figure 13C). The second ROI corresponds to a long-lasting frontal positivity in the visual modality at 1100–1600 ms that is not seen in the auditory modality (Figure 13A).

**Figure 13:**
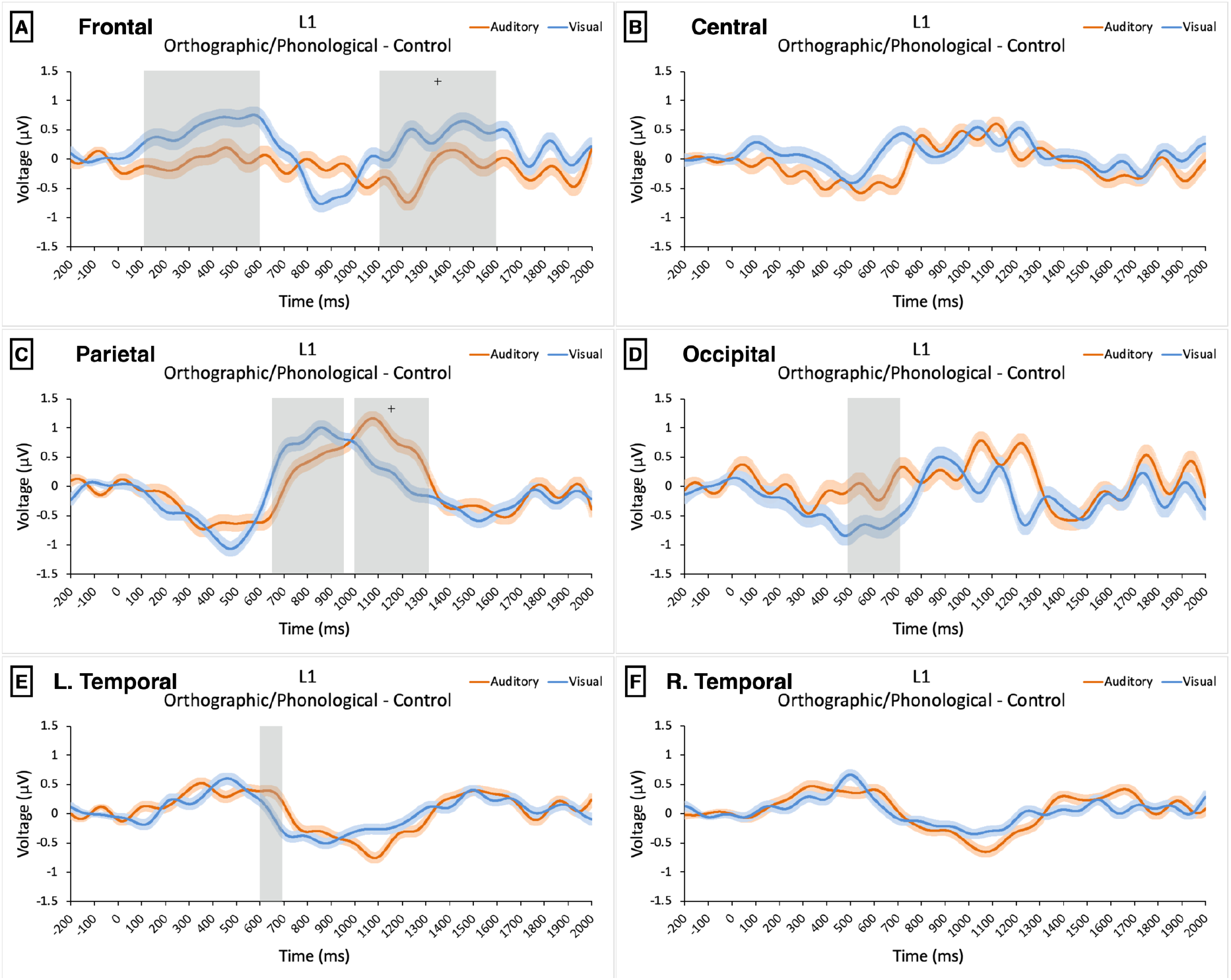
Orthographic/phonological violation difference condition ERP waveforms for each region in L1 speakers. Each waveform is the average of all electrodes in the region shown. For display purposes, waveforms are plotted with a low-pass filter at 6 Hz. Positivity is plotted upward. Orange lines represent the auditory evoked ERPs, while blue lines represent visual evoked ERPs. Standard error bars are included for each modality. The time windows of tested ROIs are highlighted in gray. ^+^p <0.10, *p < 0.05, all Holm-Bonferroni corrected.

**Figure 14.**
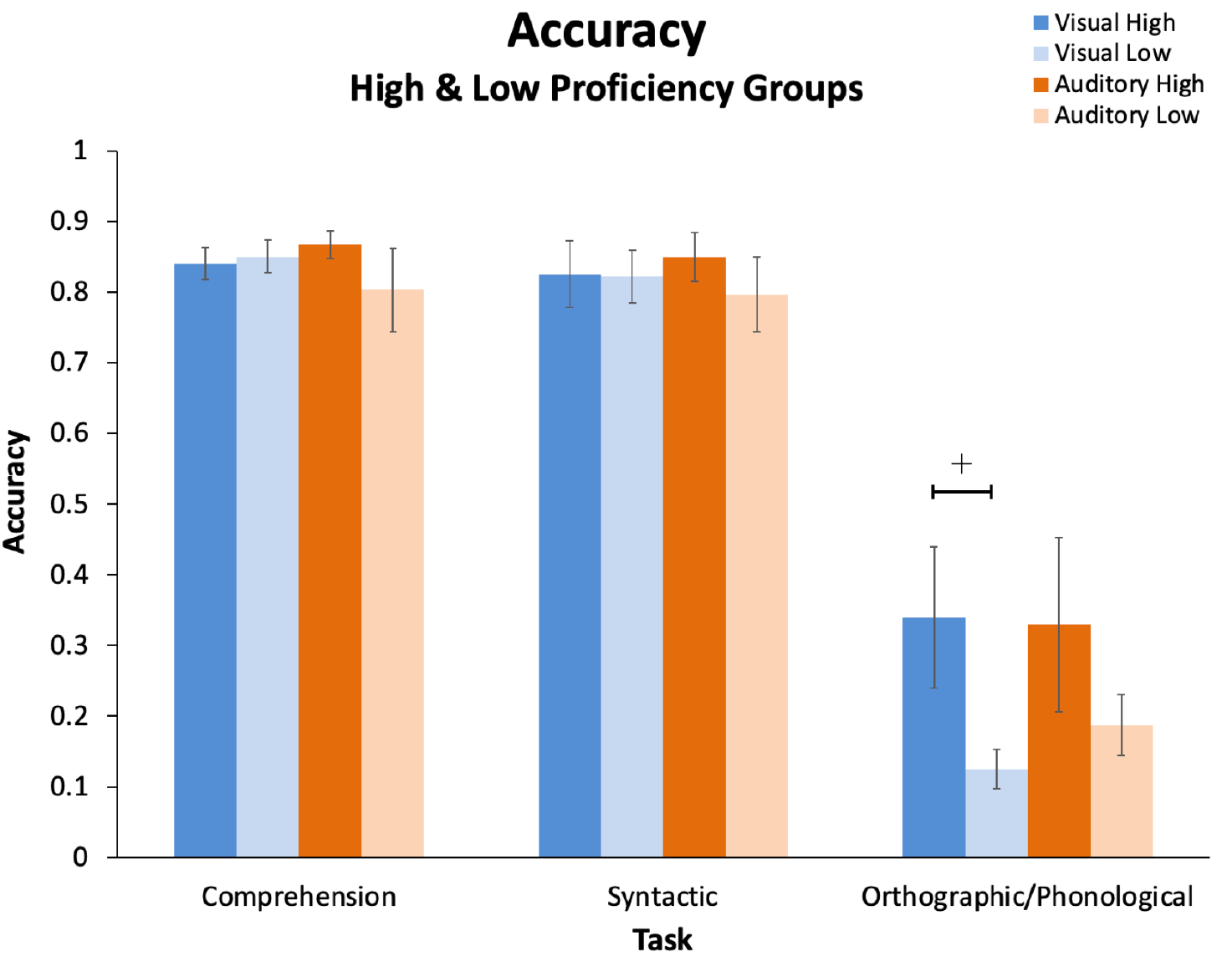
L2 accuracies in each modality (visual: blue, auditory: orange) and each proficiency group (high: dark, low: light). Error bars represent standard error. ^+^p < 0.10

The significant effects shown in Table 2 and Figure 10 represent evidence for a modality effect, demonstrating that morphosyntactic violation processing is comprised of both modality-dependent and modality-independent processing. The broader implications of this effect are discussed further in Section 4.1.

The orthographic-phonological violation difference condition elicited less statistically robust modality effects than the morphosyntactic violation difference condition. This is surprising insofar as grammatical processing is typically treated as a modality-independent process, whereas orthographic and phonological violations are by definition more closely linked to their respective modalities. Nevertheless, the effects shown in Table 3 and Figure 13, although only marginally significant, suggest the possibility of a modality effect in orthographic and phonological violation processing. However, the effect may be smaller in magnitude than that of morphosyntactic violation processing, and thus requires further investigation to clarify its characteristics. The broader implications of this effect are discussed further in Section 4.1.

### 3.2 L2 Experiment

#### 3.2.1 L2 Behavioral Results

Two-sided two-sample t-tests revealed only one marginally significant difference between the high proficiency and low proficiency groups in the orthographic/phonological judgment task, *t*(16) = 1.867, *p* = 0.0802. No other significant differences were observed between groups (within modality) or between modalities (within groups).

Similarly for response times (Figure 15), two-sided two-sample t-tests were performed to check for significant differences between groups and between modalities. When grouping all participants together, response time was significantly slower in the auditory condition for the comprehension (*t*(34) = −2.168, *p* = 0.037) and orthographic/phonological violation tasks (*t*(29) = = 2.137, *p* = 0.041). Additionally, within the high proficiency group, when comparing the orthographic judgment task with phonological judgment task, participants performed slower in the phonological judgment task with marginal significance (*t*(13) = −1.914, *p* = 0.078). Otherwise, no significant differences in response time were observed between groups or modalities.

**Figure 15.**
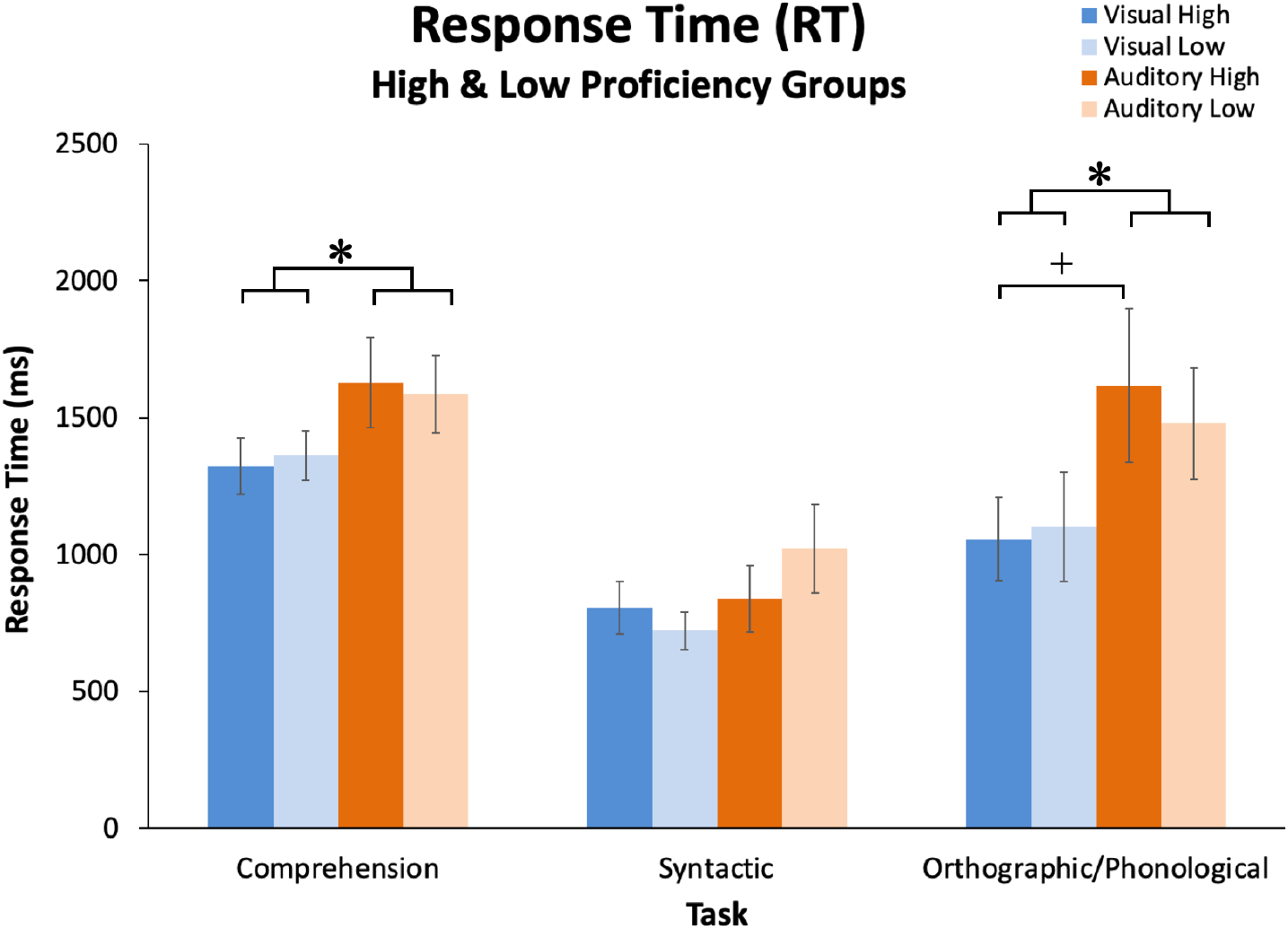
L2 response times in each modality (visual: blue, auditory: orange) and each proficiency group (high: dark, low: light). Error bars represent standard error. ^+^p < 0.10, *p < 0.05

We speculate that the slower response times in the auditory condition is either the result of internally replaying stimuli as in the phonological loop described by Baddeley and Hitch (1974) or the result of lower confidence caused by relatively little listening practice (compared to reading practice) that is typical in Japanese second language education, although there is no data comparing reading and listening exposure times in the classes attended by our participants.

#### 3.2.2 L2 Electrophysiological Results – Morphosyntactic Violations in High Proficiency L2 Learners

Single-participant data with fewer than 13 trials in any given condition remaining after artefact rejection were excluded from all conditions. Since accuracy was very low and only 10 participants had 13 or more successful trials after artefact rejection, the orthographic-phonological condition was excluded from ERP analysis in the L2 experiment. This also resulted in the exclusion of four visual datasets (one low proficiency group). An additional two auditory datasets were excluded due to experimenter error (both high proficiency group). After these exclusions, in the visual modality, there were nine high proficiency group datasets and five low proficiency group datasets, while in the auditory modality there were eight datasets in both high and low proficiency groups.

As with L1 electrophysiological data, ERPs were analyzed using difference waveforms and difference topography (i.e., the control condition subtracted from the morphosyntactic violation condition), in order to remove sensory perceptual effects that are not of interest.

Analyses of morphosyntactic violation-elicited ERPs were carried out in the same way as the L1 ERP analyses in Sections 3.1.3 and 3.1.4. We start with a qualitative examination of the whole-epoch topographical maps of voltage distribution in both modalities shown in Figure 16.

**Figure 16:**
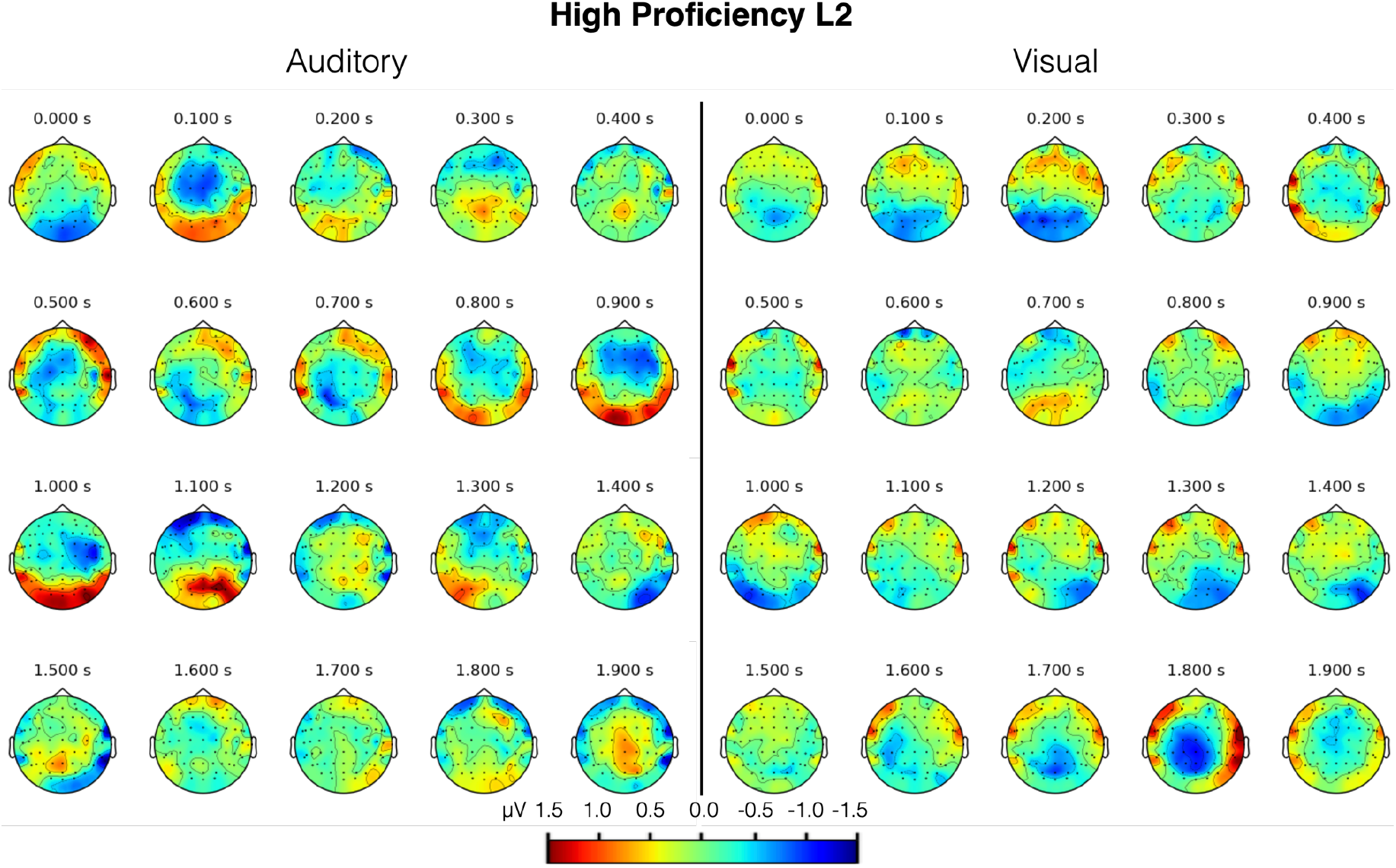
L2 high proficiency group, visual and auditory modality topographical maps of voltage distribution of the morphosyntactic violation difference condition. Each 100 ms interval is shown as a moving average with a 100 ms time window.

**Figure 17:**
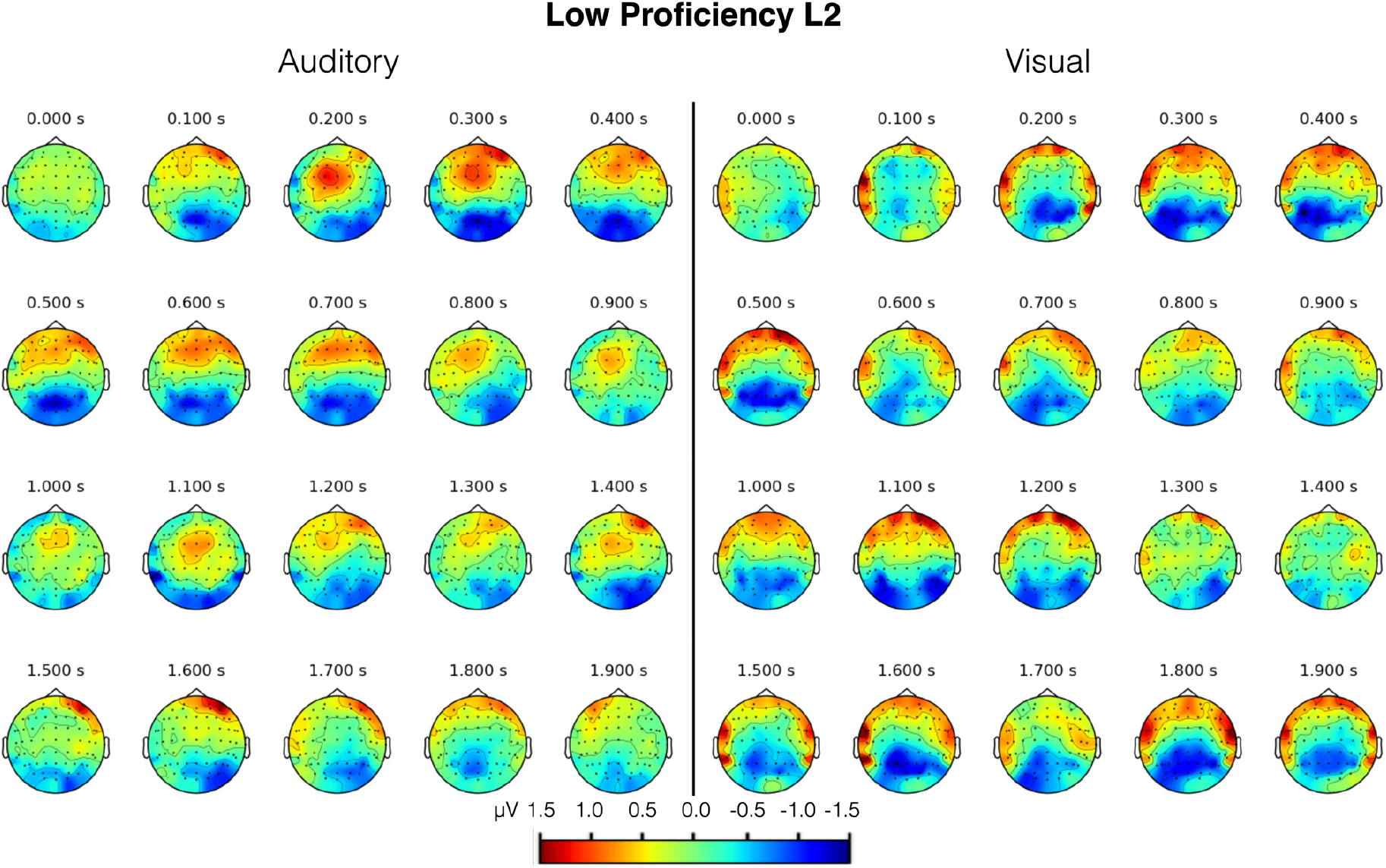
L2 low proficiency group, visual and auditory modality topographical maps of voltage distribution of the morphosyntactic violation difference condition. Each 100 ms interval is shown as a moving average with a 100 ms time window.

By visual inspection of Figure 16, a list of ROIs was compiled and tested for significance using an LME model with condition as a fixed effect and subject as a random effect (Amplitude ~ Condition + (1|Subject)). The results of these tests are shown in Table 4.

**Table 4:** P values of ROIs and their corresponding time windows in the high proficiency L2 morphosyntactic violation difference condition using the LME model Amplitude ~ Condition + (1|Subject).

| Group | Modality | Region | Time (ms) | Condition | Condition (adj.) |
| --- | --- | --- | --- | --- | --- |
| High | Auditory | Central | 100–200 | 0.151 | 0.604 |
| High | Auditory | R. Temporal | 450–550 | 0.085<sup>+</sup> | 0.425 |
| High | Auditory | Central | 800–1100 | 0.2179 | 0.4358 |
| High | Auditory | Occipital | 800–1100 | 0.196 | 0.588 |
| High | Auditory | Frontal | 1100–1300 | 0.261 | 0.261 |
| High | Visual | Occipital | 100–200 | 0.0368* | 0.1104 |
| High | Visual | Occipital | 900–1300 | 0.353 | 0.353 |
| High | Visual | L. Temporal | 1600–1900 | 0.038* | 0.076<sup>+</sup> |
| High | Visual | Parietal | 1700–1800 | 0.025* | 0.1 |
| High | Visual | R. Temporal | 1750–1850 | 0.0071** | 0.0355* |

*Adjusted p values were calculated by the Holm-Bonferroni method. <sup>+</sup> $p < 0.10$ , \* $p < 0.05$ , \*\* $p < 0.01$ , \*\*\* $p < 0.001$*

For the auditory modality in the high proficiency group, of the five ROIs tested, none had a significant main effect of condition. However, a short-lasting positivity over right temporal electrodes at 450–550 ms had a marginally significant unadjusted p value (*p* = 0.085; Figure 19F). The visual modality had more significant results. Of the five ROIs tested, four had significant unadjusted p values. After applying the Holm-Bonferroni correction, a left temporal positivity at 1600–1900 ms was marginally significant (*p* = 0.076; Figure 19E), while a shorter-lasting right temporal positivity was found to be significant (*p* = 0.0355; Figure 19F). In short, in the high proficiency L2 learner group, the auditory modality showed no significant effect of condition on ERP amplitude, while the visual modality showed a late bilateral temporal positivity.

#### 3.2.3 L2 Electrophysiological Results – Morphosyntactic Violations in Low Proficiency L2 Learners

Now turning to the low proficiency L2 learner group, the topographical map of voltage distribution for the morphosyntactic violation difference condition is shown for both modalities in Figure 17.

The topographical maps of Figure 17 suggest that both modalities are marked primarily by long-lasting centrofrontal positivities and parietooccipital negativities, with the visual modality being more bilaterally temporal in its distribution. By visual inspection of Figure 17, a list of ROIs was compiled and tested for significance using an LME model with condition as a fixed effect and subject as a random effect (Amplitude ~ Condition + (1|Subject)). The results of these tests are shown in Table 5.

**Table 5:** P values of ROIs and their corresponding time windows in the low proficiency L2 morphosyntactic violation difference condition using the LME model Amplitude ~ Condition + (1|Subject).

| Group | Modality | ROI | Time (ms) | Condition | Condition (adj.) |
| --- | --- | --- | --- | --- | --- |
| Low | Auditory | Occipital | 100–800 | 0.00671** | 0.02684* |
| Low | Auditory | Parietal | 100–800 | 0.0344* | 0.1032 |
| Low | Auditory | Central | 200–400 | 0.091142<sup>+</sup> | 0.091142<sup>+</sup> |
| Low | Auditory | Frontal | 300–700 | 0.00105** | 0.0063** |
| Low | Auditory | Frontal | 1200–1800 | 0.00644** | 0.0322* |
| Low | Auditory | Occipital | 1100–1700 | 0.0465* | 0.093<sup>+</sup> |
| Low | Visual | L. Temporal | 100–700 | 0.07416<sup>+</sup> | 0.29664 |
| Low | Visual | R. Temporal | 200–700 | 0.00854** | 0.05124<sup>+</sup> |
| Low | Visual | Parietal | 200–800 | 0.087<sup>+</sup> | 0.261 |
| Low | Visual | Occipital | 1000–1200 | 0.0507<sup>+</sup> | 0.2535 |
| Low | Visual | Frontal | 1000–1100 | 0.1567 | 0.3134 |
| Low | Visual | Parietal | 1500–1900 | 0.2512 | 0.2512 |

Adjusted *p* values were calculated by the Holm-Bonferroni method. <sup>+</sup>*p* < 0.10, \**p* < 0.05, \*\**p* < 0.01, \*\*\**p* < 0.001

As can be seen in Table 5, of the 6 ROIs tested in the auditory modality 3 ROIs remained significant after applying the Holm-Bonferroni correction procedure, while 2 were marginally significant. Auditory morphosyntactic violation processing was marked by an occipitoparietal negativity at 100–800 ms (occipital *p* = 0.02684; parietal *p* = 0.1032), during which a centrofrontal positivity arose, (central at 200–400 ms, *p* = 0.091142; frontal at 300–700 ms, *p* = 0.0063). Finally, a similar frontal positivity arose at 1200–1800 ms (*p* = 0.0322) and an occipital negativity arose at 1100–1700 ms (*p* = 0.093).

Of the 6 ROIs tested in the visual modality, prior to Holm-Bonferroni correction, 1 ROI was significant and 3 were marginally significant. After correction, only 1 ROI was marginally significant, a right temporal positivity at 200–700 ms (*p* = 0.05124). Despite a visual comparison between the modalities suggesting similar amplitudes and distributions, it is unclear why fewer regions reached significance in the visual modality. One contributing factor may be that topographic distributions straddled multiple regions thereby distributing the analysis across separate ROIs, weakening statistical power. Nevertheless, we qualitatively comment on the topographical maps (Figure 17) and ERP waveforms (Figure 20) in the Discussion section below.

#### 3.2.4 L2 Electrophysiological Results – The Modality Effect in High and Low Proficiency L2 Learners

Although some discussion of modality-specific differences has been given for each L2 group in Sections 3.2.2 and 3.2.3, we now analyze the L2 modality effect with the same procedure implemented on L1 speakers in Section 3.1. We first take the morphosyntactic violation difference condition in each modality and subtract the visual difference condition from the auditory difference condition, giving us the modality difference condition for each L2 proficiency group, as shown in Figure 18.

**Figure 18:**
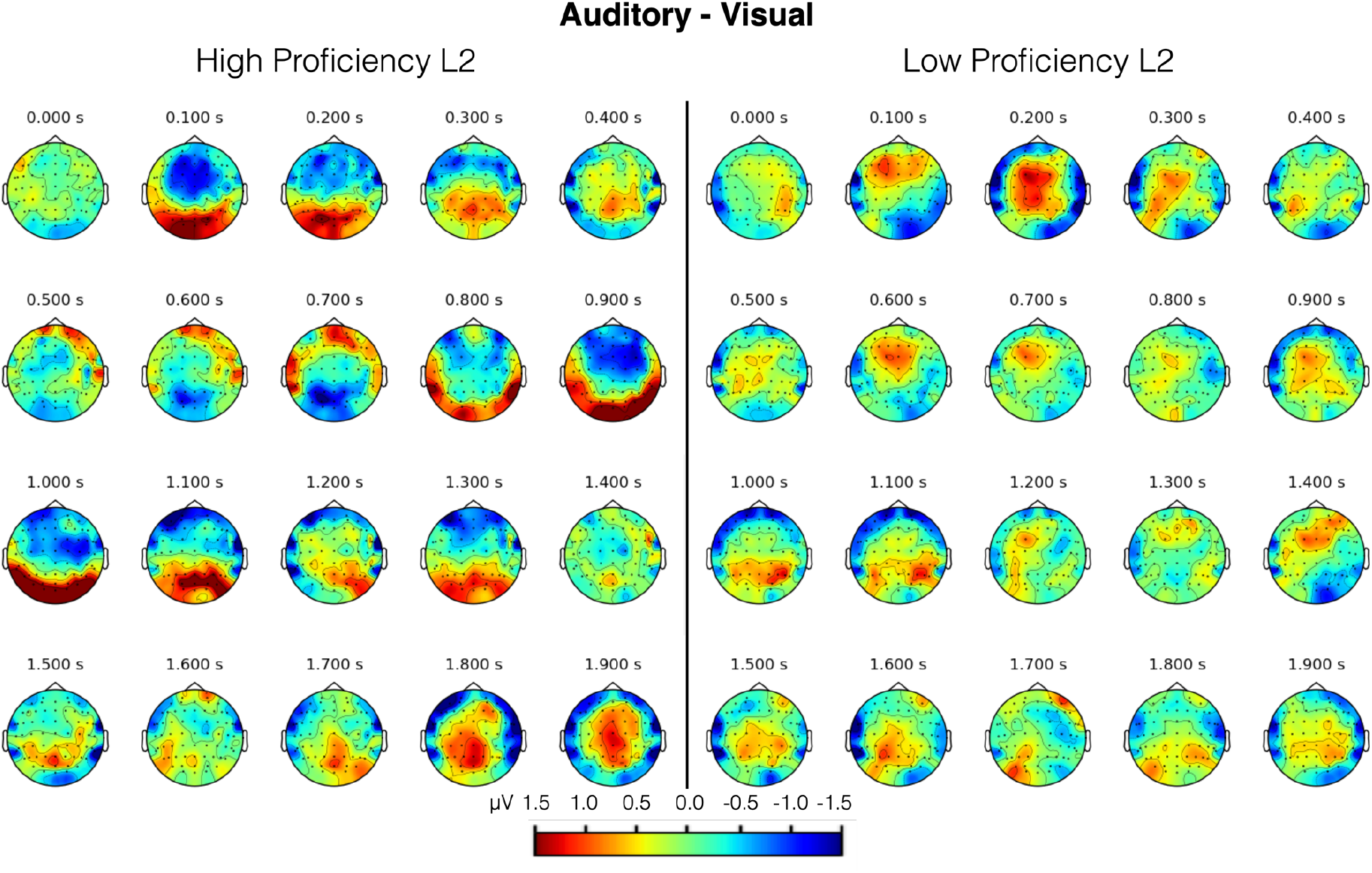
Topographical map of voltage distribution of the modality effect in L2 morphosyntactic violation processing across time. Each 100 ms interval is shown as a moving average with a 100 ms time window. Voltage was computed using the morphosyntactic violation difference condition of each modality and subtracting the visual from the auditory.

Upon visual inspection of the topographical maps of Figure 18, it is readily apparent that high proficiency L2 learners have greater modality-specific differences when processing morphosyntactic violations. As done with native speakers in Section 3.1, we compile a list of ROIs from Figure 18 (Table 6) and test their significance using an LME model with modality and condition as fixed effects and subject as a random effect (Amplitude ~ Modality * Condition + (1|Subject)). As described in Section 3.1.3, a significant main effect of modality or condition does not correspond to a modality effect, rather to (potentially) perceptual processes or condition-specific processes. We instead look for the interaction between modality and condition to indicate a modality effect.

**Table 6:** P values of ROIs and their corresponding time windows in the L2 morphosyntactic violation difference condition using the LME model Amplitude ~ Modality * Condition + (1|Subject).

| Group | ROI | Time (ms) | Modality | Condition | Modality:<br>Condition | Modality:<br>Condition<br>(adj.) |
| --- | --- | --- | --- | --- | --- | --- |
| High | Central | 100–200 | 0.401 | 0.402 | 0.122 | 0.366 |
| High | Occipital | 100–200 | 0.0854<sup>+</sup> | 0.0456* | 0.0761<sup>+</sup> | 0.4566 |
| High | Parietal | 200–400 | 5.64e-06*** | 0.135 | 0.0415* | 0.3735 |
| High | Occipital | 500–700 | 0.5657 | 0.2311 | 0.1701 | 0.3402 |
| High | Frontal | 600–700 | 0.8081 | 0.1422 | 0.0273* | 0.3003 |
| High | Central | 800–1000 | 0.7701 | 0.4491 | 0.0679<sup>+</sup> | 0.4753 |
| High | Occipital | 800–1100 | 0.813 | 0.2915 | 0.0447* | 0.3576 |
| High | Frontal | 900–1300 | 0.0399* | 0.45063 | 0.24085 | 0.24085 |
| High | Parietal | 1700–1900 | 2.68e-05*** | 0.0517<sup>+</sup> | 0.0932<sup>+</sup> | 0.466 |
| High | L. Temporal | 1800–1900 | 1.98e-04*** | 0.103685 | 0.096109<sup>+</sup> | 0.384436 |
| High | R. Temporal | 1800–1900 | 6.12e-05*** | 0.0348* | 0.032* | 0.32 |
| Low | Central | 100–200 | 0.4688 | 0.1102 | 0.0262* | 0.2096 |
| Low | Central | 600–700 | 0.00974** | 0.14539 | 0.07374<sup>+</sup> | 0.51618 |
| Low | L. Temporal | 1500–1600 | 0.000676*** | 0.289764 | 0.286619 | 1.0 |
| Low | L. Temporal | 900–1100 | 0.02251* | 0.65105 | 0.3478 | 1.0 |
| Low | Parietal | 900–1100 | 0.0793<sup>+</sup> | 0.4035 | 0.4084 | 1.0 |
| Low | L. Temporal | 200–300 | 0.00866** | 0.10252 | 0.41393 | 1.0 |
| Low | Parietal | 1500–1600 | 0.005699** | 0.160507 | 0.5078 | 1.0 |
| Low | R. Temporal | 1800–1900 | 0.148 | 0.155 | 0.553 | 0.553 |

*Adjusted p values were calculated by the Holm-Bonferroni method. Adjusted p values above 1.0 were reduced to 1.0* <sup>+</sup>*p* < 0.10, \**p* < 0.05, \*\**p* < 0.01, \*\*\**p* < 0.001

In the high proficiency group, we identified 11 ROIs based on the topographical maps in Figure 18. Of the 11 ROIs, 4 unadjusted p values were significant and 4 were marginally significant (Table 6, Figure 19). After applying the Holm-Bonferroni correction procedure, none of the ROIs were significant. This is likely due to a combination of factors, namely a small sample size, a small effect, and the inclusion of many tests (which increases the factor for p value adjustment). In the low proficiency group, we identified 8 ROIs from Figure 18. Of the 8 ROIs, 1 unadjusted p value was significant and 1 was marginally significant (Table 6, Figure 20). After applying the correction procedure, none of the ROIs were found to be significant. The same statistical challenges affecting the high proficiency group (sample size, effect size, many tests) affect the low proficiency group as well.

**Figure 19:**
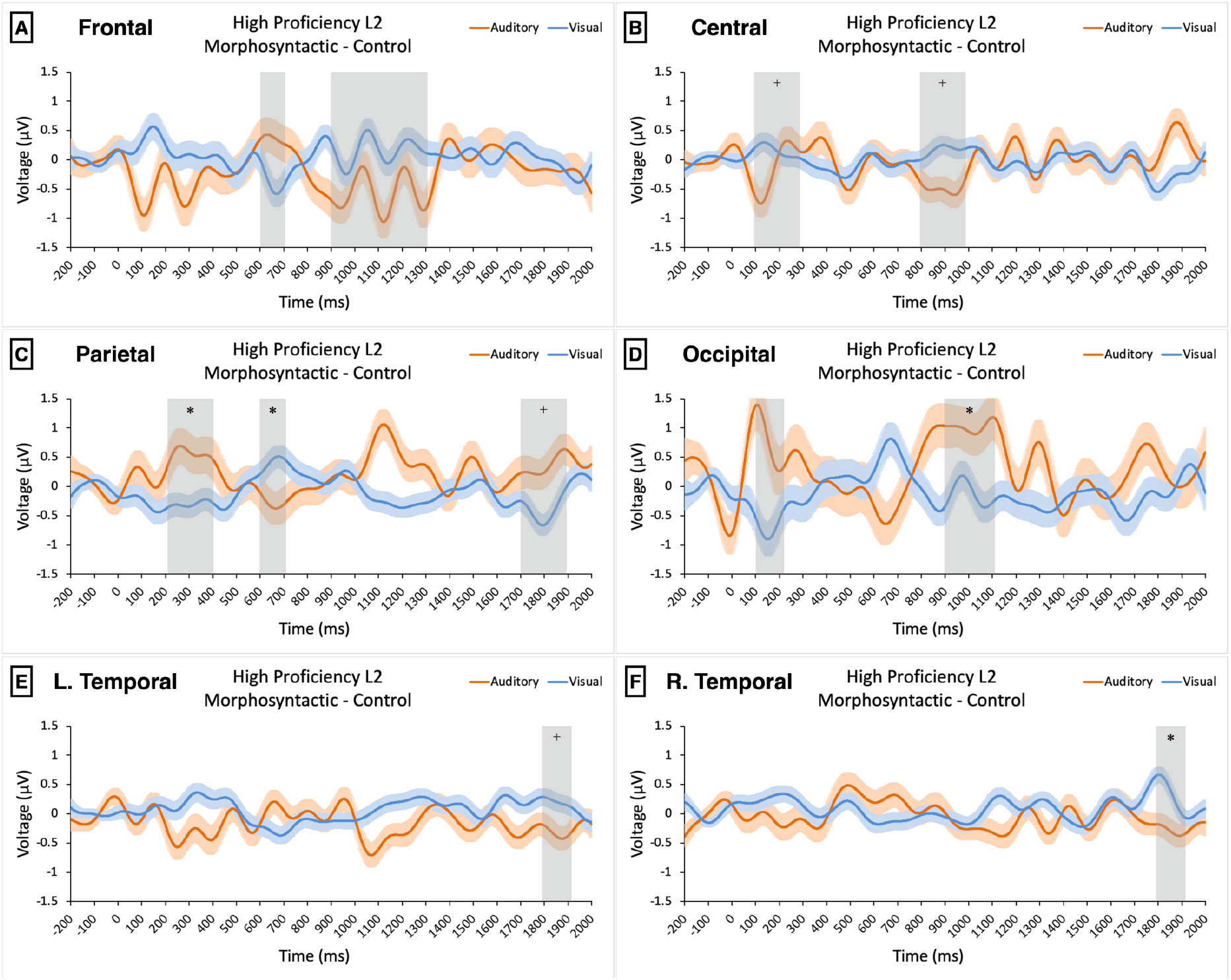
Morphosyntactic violation difference condition ERP waveforms for each region in high proficiency L2 learners. Each waveform is the average of all electrodes in the region shown. For display purposes, waveforms are plotted with a low-pass filter at 6 Hz. Orange lines represent the auditory evoked ERPs, while blue lines represent visual evoked ERPs. Standard error bars are included for each modality. The time windows of tested ROIs are highlighted in gray. ^+^p <0.10, *p < 0.05, all unadjusted.

**Figure 20:**
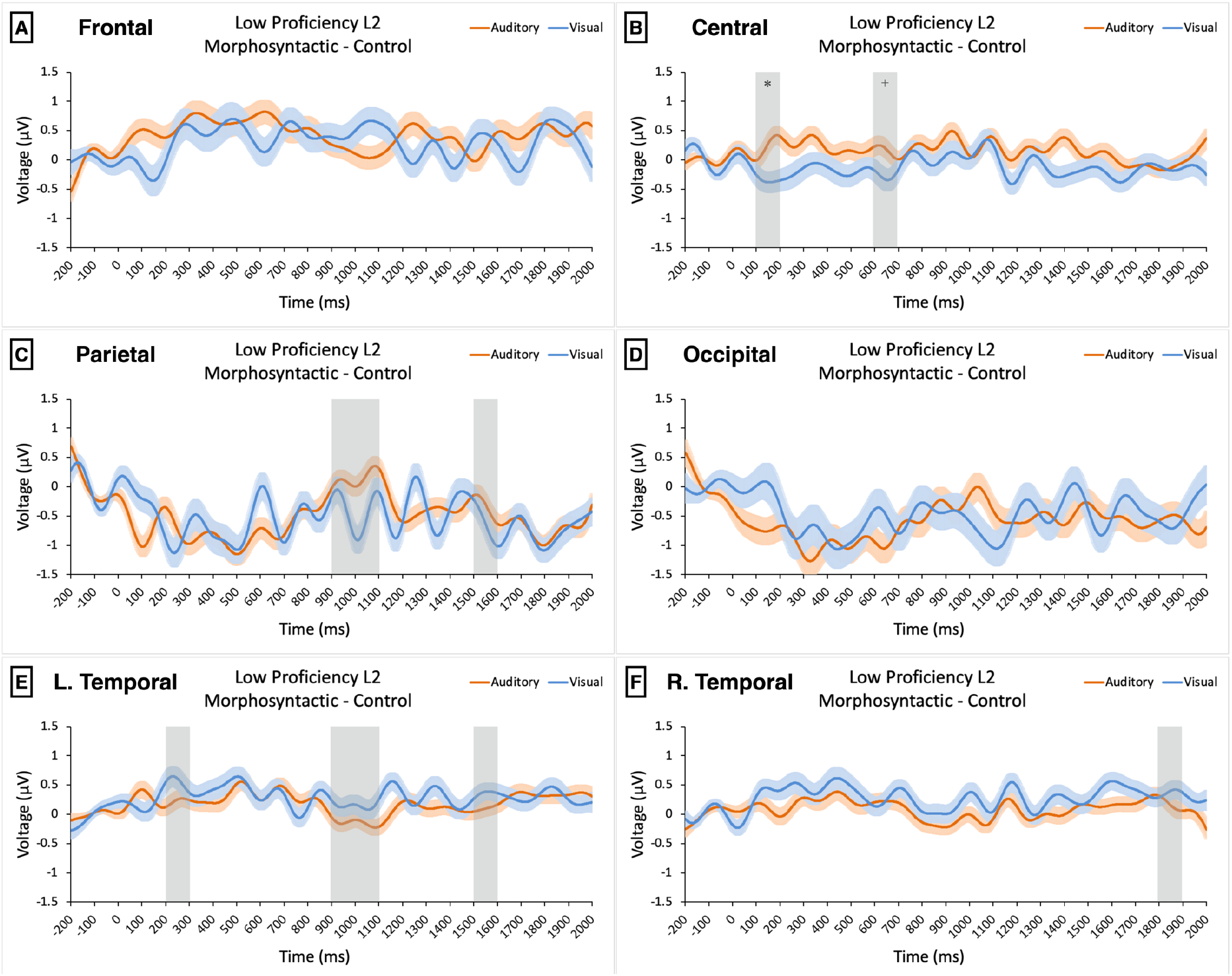
Morphosyntactic violation difference condition ERP waveforms for each region in low proficiency L2 learners. Each waveform is the average of all electrodes in the region shown. For display purposes, waveforms are plotted with a low-pass filter at 6 Hz. Orange lines represent the auditory evoked ERPs, while blue lines represent visual evoked ERPs. Standard error bars are included for each modality. The time windows of tested ROIs are highlighted in gray. ^+^p <0.10, *p < 0.05, all unadjusted.

Although it is difficult to precisely characterize the nature of the modality effect in L2 learners, we can qualitatively state that there is a greater effect of modality on language processing in higher proficiency learners than in lower proficiency learners, as evidenced by the topographical maps of Figure 18 and the larger number of unadjusted significant p values in high proficiency learners (4 significant, 4 marginally significant) compared to low proficiency learners (1 significant, 1 marginally significant). This separation of language processing streams is qualitatively similar to native speakers, although the modality effect in high proficiency learners seems to share few characteristics with that of native speakers.

#### 3.2.4 L2 Electrophysiological Results – Nativization by Modality in L2 Learners

We next consider differences between L1 speakers and L2 learners. Continuing with the same procedure used heretofore, we begin with a topographical map of voltage distribution computed by subtracting the morphosyntactic difference condition of high proficiency L2 learners from that of L1 speakers, and we do this for both modalities, as shown in Figure 21. We do the same with low proficiency L2 learners, as shown in Figure 22.

**Figure 21:**
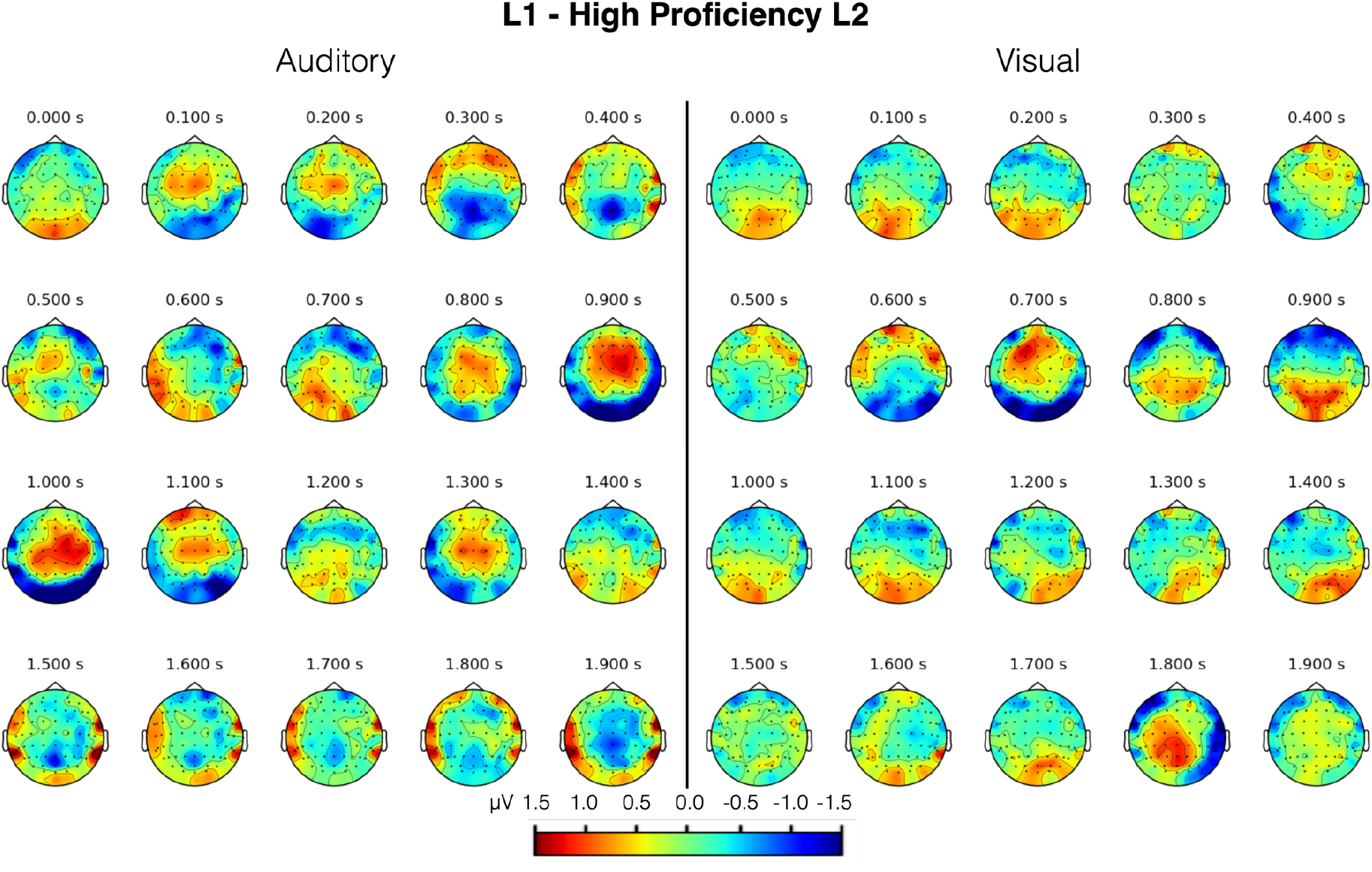
Topographical map of voltage distribution for L1 morphosyntactic violation difference condition minus the high proficiency L2 morphosyntactic violation difference condition, for each modality. Each 100 ms interval is shown as a moving average with a 100 ms time window.

**Figure 22:**
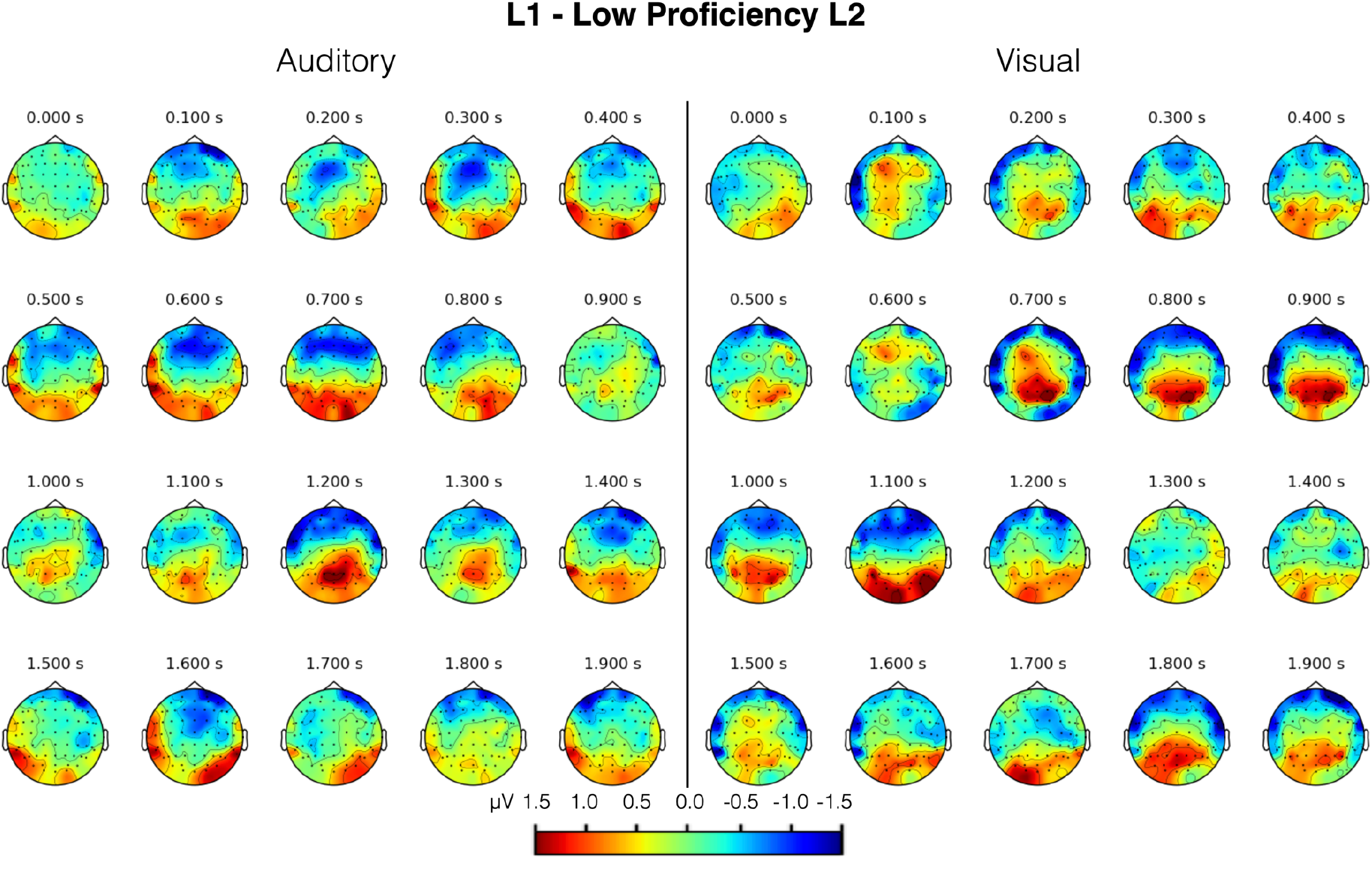
Topographical map of voltage distribution for L1 morphosyntactic violation difference condition minus the low proficiency L2 morphosyntactic violation difference condition, for each modality. Each 100 ms interval is shown as a moving average with a 100 ms time window. Voltage was computed using the morphosyntactic violation difference condition of each modality and subtracting the visual from the auditory.

From Figures 21 and 22, a list of ROIs was compiled for each proficiency group in each modality and tested for significance using an LME model with fixed effects of condition and group (high or low proficiency L2 learners and L1 speakers) and a random effect of subject (Amplitude ~ Condition * Group + (1|Subject)), shown in Table 7.

**Table 7:**
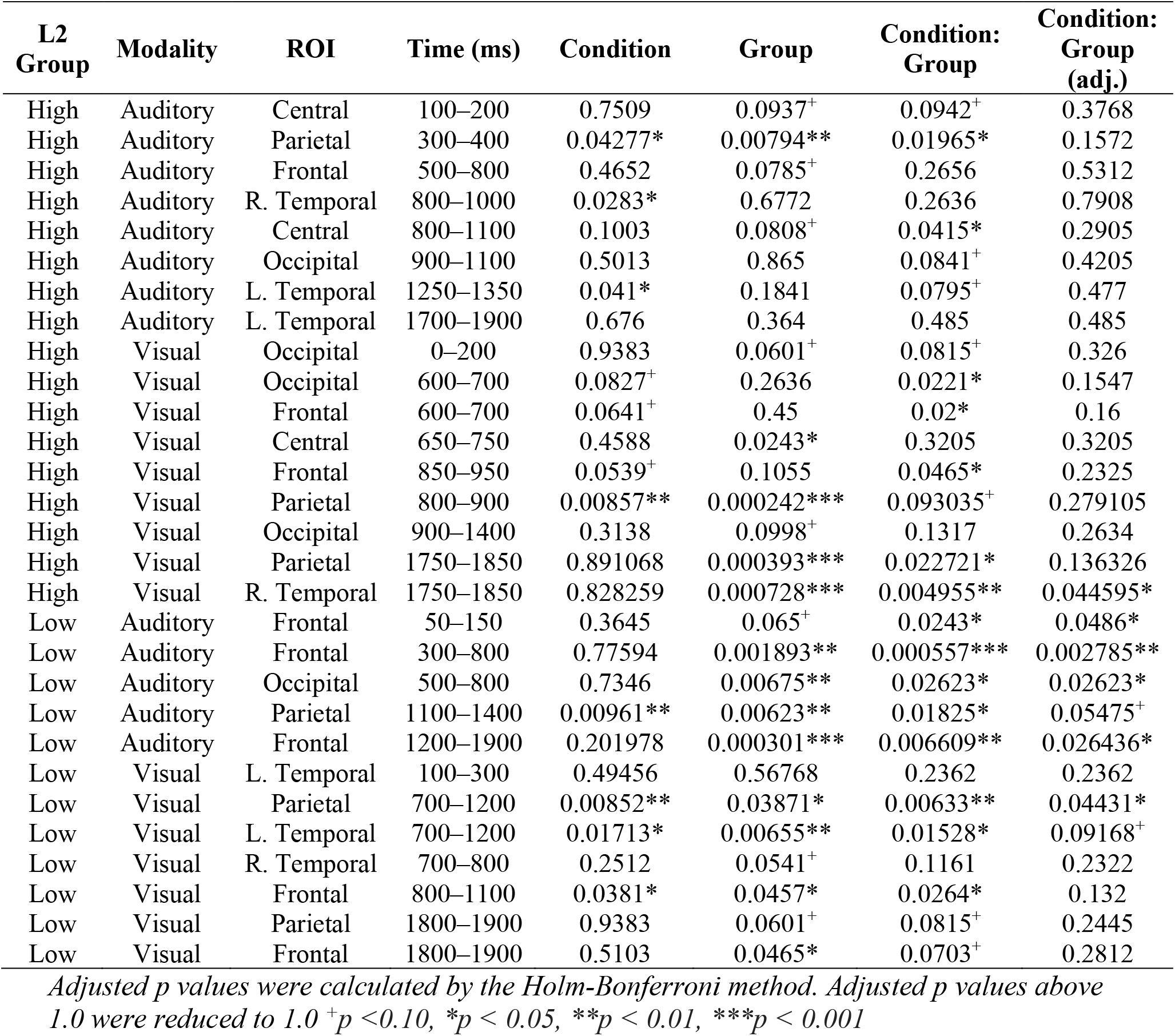
P values of ROIs and their corresponding time windows in the morphosyntactic violation difference condition of L2 learners compared to L1 speakers using the LME model Amplitude ~ Condition * Group + (1|Subject).

| L2 Group | Modality | ROI | Time (ms) | Condition | Group | Condition: Group | Condition: Group (adj.) |
| --- | --- | --- | --- | --- | --- | --- | --- |
| High | Auditory | Central | 100–200 | 0.7509 | 0.0937<sup>+</sup> | 0.0942<sup>+</sup> | 0.3768 |
| High | Auditory | Parietal | 300–400 | 0.04277* | 0.00794** | 0.01965* | 0.1572 |
| High | Auditory | Frontal | 500–800 | 0.4652 | 0.0785<sup>+</sup> | 0.2656 | 0.5312 |
| High | Auditory | R. Temporal | 800–1000 | 0.0283* | 0.6772 | 0.2636 | 0.7908 |
| High | Auditory | Central | 800–1100 | 0.1003 | 0.0808<sup>+</sup> | 0.0415* | 0.2905 |
| High | Auditory | Occipital | 900–1100 | 0.5013 | 0.865 | 0.0841<sup>+</sup> | 0.4205 |
| High | Auditory | L. Temporal | 1250–1350 | 0.041* | 0.1841 | 0.0795<sup>+</sup> | 0.477 |
| High | Auditory | L. Temporal | 1700–1900 | 0.676 | 0.364 | 0.485 | 0.485 |
| High | Visual | Occipital | 0–200 | 0.9383 | 0.0601<sup>+</sup> | 0.0815<sup>+</sup> | 0.326 |
| High | Visual | Occipital | 600–700 | 0.0827<sup>+</sup> | 0.2636 | 0.0221* | 0.1547 |
| High | Visual | Frontal | 600–700 | 0.0641<sup>+</sup> | 0.45 | 0.02* | 0.16 |
| High | Visual | Central | 650–750 | 0.4588 | 0.0243* | 0.3205 | 0.3205 |
| High | Visual | Frontal | 850–950 | 0.0539<sup>+</sup> | 0.1055 | 0.0465* | 0.2325 |
| High | Visual | Parietal | 800–900 | 0.00857** | 0.000242*** | 0.093035<sup>+</sup> | 0.279105 |
| High | Visual | Occipital | 900–1400 | 0.3138 | 0.0998<sup>+</sup> | 0.1317 | 0.2634 |
| High | Visual | Parietal | 1750–1850 | 0.891068 | 0.000393*** | 0.022721* | 0.136326 |
| High | Visual | R. Temporal | 1750–1850 | 0.828259 | 0.000728*** | 0.004955** | 0.044595* |
| Low | Auditory | Frontal | 50–150 | 0.3645 | 0.065<sup>+</sup> | 0.0243* | 0.0486* |
| Low | Auditory | Frontal | 300–800 | 0.77594 | 0.001893** | 0.000557*** | 0.002785** |
| Low | Auditory | Occipital | 500–800 | 0.7346 | 0.00675** | 0.02623* | 0.02623* |
| Low | Auditory | Parietal | 1100–1400 | 0.00961** | 0.00623** | 0.01825* | 0.05475<sup>+</sup> |
| Low | Auditory | Frontal | 1200–1900 | 0.201978 | 0.000301*** | 0.006609** | 0.026436* |
| Low | Visual | L. Temporal | 100–300 | 0.49456 | 0.56768 | 0.2362 | 0.2362 |
| Low | Visual | Parietal | 700–1200 | 0.00852** | 0.03871* | 0.00633** | 0.04431* |
| Low | Visual | L. Temporal | 700–1200 | 0.01713* | 0.00655** | 0.01528* | 0.09168<sup>+</sup> |
| Low | Visual | R. Temporal | 700–800 | 0.2512 | 0.0541<sup>+</sup> | 0.1161 | 0.2322 |
| Low | Visual | Frontal | 800–1100 | 0.0381* | 0.0457* | 0.0264* | 0.132 |
| Low | Visual | Parietal | 1800–1900 | 0.9383 | 0.0601<sup>+</sup> | 0.0815<sup>+</sup> | 0.2445 |
| Low | Visual | Frontal | 1800–1900 | 0.5103 | 0.0465* | 0.0703<sup>+</sup> | 0.2812 |

*Adjusted p values were calculated by the Holm-Bonferroni method. Adjusted p values above 1.0 were reduced to 1.0* <sup>+</sup>*p* < 0.10, \**p* < 0.05, \*\**p* < 0.01, \*\*\**p* < 0.001

Taking either comparison by itself is not very informative, but by comparing the difference between L1 and high proficiency L2 speakers with the difference between L1 and low proficiency L2 speakers, we can form a rough idea of how language processing changes with acquired proficiency.

In particular, by focusing on Figures 21 and 22, we readily see that low proficiency L2 learners differ from L1 speakers to a greater degree than high proficiency L2 learners do. Table 7 supports this assertion by revealing more and longer-lasting significant differences for the low proficiency group across both modalities. Specifically, in the auditory modality, the high proficiency group had no significant ROIs post p value correction, while the low proficiency group had 4 (with an average duration of 400 ms). In the auditory modality, the high proficiency group had 1 significant ROI (duration of 100 ms), while the low proficiency group had 1 significant ROI (duration of 500 ms) and 1 marginally significant ROI (duration of 500 ms). Taken together, the evidence suggests that with improved L2 proficiency comes more nativelike language processing, a process that could be called “(neurophysiological) nativization.” However, since not all levels of proficiency are represented in our participant pool and since participants are grouped together, this does not preclude the possibility that some high-level L2 speakers employ non-nativelike strategies in L2 language processing.

When comparing the degree of nativization in each modality, an interesting discrepancy becomes apparent. Namely, the lower-proficiency group had 4 significant ROIs in the auditory modality and only 1 in the visual modality. This could be interpreted in one of two ways, either that L2 auditory language processing starts out more distinct from native auditory language processing than visual language processing, or that some degree of nativization has already taken place in the visual modality even among low-proficiency L2 learners. In the former case, it would not be surprising that L2 learners who are a) very familiar with an orthography like the Latin alphabet used in Spanish and English and b) primarily study in the written format have less difficulty processing written language than spoken language, which may contain unfamiliar sounds and difficult to understand native pronunciation (which may be quite different from the pronunciation of the subvocalization non-native learners use while reading).

In stark contrast, among high-proficiency L2 learners, the auditory modality had no significant ROIs, while the visual modality had 1 significant ROI. The number of significant ROIs in the visual modality did not decrease with proficiency, while the number of significant ROIs in the auditory modality did drop dramatically. What exactly this means is certainly subject to debate, but it may suggest that higher-proficiency learners have had more effective neurophysiological nativization in the auditory modality than in the visual modality. This is further discussed in Section 4.2 below.

## 4 Discussion

A multi-modal ERP experiment was conducted to establish whether there exists a neurophysiological modality effect in language processing that is not explicable by simple perceptual differences. The experiment was conducted with both native and non-native speakers, who were divided into two proficiency groups (high and low).

### 4.1 The Modality Effect in Native Language Processing

In the native speaker group, very few behavioral differences were observed across modalities. Specifically, native speakers more accurately judged orthographic violations (misspellings) than phonological violations (mispronunciations). The lower accuracy in the auditory modality for the orthographic/phonological judgment task is likely due to two factors. First, the featurally underspecified lexicon (FUL) model predicts that some phonological features are underspecified compared to others (Lahiri & Reetz, 2002). This phenomenon is easily understood in English by considering that changing ‘rainbow’ ([+coronal]) to ‘raimbow’ ([-coronal]) is more acceptable and less jarring than the same change in the opposite direction, as in ‘tame tiger’ ([-coronal]) to ‘tane tiger’ ([+coronal]). Violations of such underspecified phonemes are more likely to go unnoticed, even by native speakers. Second, because participants came from different Spanish-speaking countries, dialectical differences may account for some participants failing to notice mispronunciations where others successfully spotted the error.

Aside from orthographic/phonological task accuracy, no significant differences were observed in the behavioral results. Because the stimuli were designed to be accessible to non-native speakers, the tasks were very easy for the native speakers in this experiment, giving rise to a ceiling effect. Therefore, we cannot confidently assert one way or the other whether accuracies or response times of the comprehension and morphosyntactic judgment tasks would have had a significant effect of modality with more difficult stimuli.

The primary interest of this study, however, lies in the electrophysiological results. By analyzing the difference conditions, whereby the control condition was subtracted from the violation conditions, we can safely regard modality-specific differences as a modality effect that is not an effect of sensory perception, but rather an effect of distinct language processing streams across modalities.

We observed various ERP components exhibiting modality-specific features. Perhaps the most important observation made herein pertains to the most commonly studied ERP components, the N400 and P600. One prominent interpretation is that the N400 indexes retrieval and the P600 indexes integration (Aurnhammer, Delogu, Schulz, Brouwer, & Crocker, 2021; Brouwer & Crocker, 2017; Delogu, Brouwer, & Crocker, 2019). In our experiment, in both the morphosyntactic violation condition and the orthographic/phonological violation condition, both modalities showed both of these components. However, while N400 characteristics were entirely consistent across modalities, P600 characteristics were modality-dependent. Specifically, although onset latencies were consistent across modalities, the auditory P600 had a distinctly more gradual slope and later peak latency (peaking about 350 ms later). While the auditory P600 was peaking, the visual P600 had already diminished nearly to zero (1100–1300 ms, Figure 10C).

The consistency of the N400 across modalities suggests that the lexical retrieval indexed by the N400 is taking place post orthographic or phonological mapping on to semantic representations, otherwise, we would have expected to see modality-specific features in the N400. The consistency of the N400 latency across modalities in both the morphosyntactic violation condition and orthographic/phonological violation condition serves as further evidence that using critical syllable onset to align auditory ERPs is a valid approach, and that the more gradual slope of the P600 in the auditory modality is not the result of misaligned ERPs.

By contrast, the presence of this modality effect in the P600 indicates that the integration thought to be indexed by the P600 must rely on modality-specific processes. These modality-specific processes may consist of a backpropagation for probability weight updating, as many studies have suggested the P600 is primarily related to expectancy (Aurnhammer et al., 2021; Fitz & Chang, 2019), or alternatively, may be related to monitoring processes (Hagoort, 2003; van Herten, Kolk, & Chwilla, 2005). Whether due to backpropagation, monitoring, or something else altogether (e.g., sensory-input related recall), it is clear from our results that the language processing indexed by the P600 must go down to the level of orthographic or phonological representation.

Aside from the N400 and P600, in the morphosyntactic violation condition, we observed a negativity over occipital electrodes at 600–700 ms in the visual modality that was absent in the auditory modality (Figure 10D), and a left temporal negativity at 1100–1200 ms in the auditory modality that was absent in the visual modality (Figure 10E). As previously noted, the visual evoked occipital negativity and the auditory evoked left temporal negativity were topographically distributed over the primary visual cortex and the primary auditory cortex, respectively. Although EEG has notoriously poor spatial resolution, these distributions may correspond to modality-specific processes taking place in their respective sensory perception hubs.

One notable difference between the orthographic/phonological violation condition and the morphosyntactic violation condition ERPs was the magnitude of the N400, where the former elicits much greater negativities (in both modalities) than the latter. Since the N400 is associated with lexical retrieval (Delogu et al., 2019), this may index an increased difficulty of retrieval when a word is misspelled or mispronounced compared to incorrectly inflected, which is more of a problem of integration rather than retrieval. Aside from this difference, the orthographic/phonological violation condition was also marked by a frontal positivity in the visual modality at 1100–1600 ms.

Returning to the question of language processing streams from the perspective of modality-specificity (Figure 1), our results readily refute the possibility of independent streams model (as evidenced by the sweeping similarities across modalities, including the N400) as well as a merging model that does not split (as evidenced by modality-specific features of the P600). We can also rule out a merging and splitting streams model (Figure 1 Model 3), since visual and auditory evoked ERPs were largely indistinguishable in the later stages of our epoched data. That leaves interactive/parallel models, which either end with amodal language processing or modality-specific language processing (Figure 1 Models 4 and 5, respectively). Since it is unclear exactly when language processing of a given stimulus ends, and since we observed some differences in the morphosyntactic violation condition and the orthographic/phonological violation condition, we cannot specify whether the final stages of language processing are modality-independent (Figure 1 Model 4) or modality-dependent (Figure 1 Model 5). But taken together, our results provide evidence for interactive/parallel processing between amodal language processing and modality-specific language processing.

Our finding of a neurophysiological modality effect in native language processing contradicts the results of Balconi & Pozzoli (2005), which showed that there was no effect of stimulus modality on either the N400 or P600, and extends on the results of Holcomb, et al. (1992) and Osterhout & Holcomb (1993), who described some minor variations between modalities.

In particular, our methodology stands apart from the previous studies in two important ways. First, we have aligned auditory ERPs to critical syllable onset, removing temporal discrepancies due to stimulus features rather than differences in language processing. Second, we have used the difference condition of each modality to make comparisons. By not using the difference condition, results are confounded by non-(morpho)syntactic processing differences such as sensory perception. In fact, upon closer inspection of the figures and data tables of Balconi & Pozzoli (2005), it is clear that the congruous sentence condition in their experiment elicited unique ERP characteristics in each modality, and after subtracting the congruous condition from each modality, the auditory P600 peak latency was more than a standard deviation later than the visual P600 peak latency. We therefore attribute some of the difference in our conclusions to a difference in methodological approach.

### 4.2 The Modality Effect in L2 Learners

In agreement with previous studies, the high proficiency L2 group showed greater similarities to L1 speakers than the low proficiency group in both modalities.

Low-proficiency L2 learners exhibited long-lasting frontocentral positivities coupled by occipitoparietal positivities in both modalities, although the visual modality had more bilateral temporal positivities than the auditory modality. The high proficiency L2 group had markedly more modality-specific differences in the morphosyntactic violation difference condition. Coupling this with the results that high proficiency L2 learners showed greater similarities to L1 speakers and that L1 speakers exhibited a strong modality effect, we can confidently assert that with increased proficiency comes increased modality-specific specialization in language processing.

Regarding modality-specific nativization, in the low-proficiency L2 group, we observed greater similarities between L2 learners and L1 speakers in the visual modality than in the auditory modality. This effect could be due to a combination of familiarity with the Latin script used in Spanish and an unfamiliarity with native Spanish pronunciation, thereby putting auditory language processing at a distinct disadvantage in lower-proficiency learners. On the other hand, since all low-proficiency L2 learners in our experiment had already completed at least one semester of Spanish instruction, even these low-proficiency L2 learners may have already nativized to some extent in the visual modality prior to participation in our experiment. Norms of foreign language education in Japan (for example, the practice of writing foreign words in katakana, a Japanese syllabary which cannot adequately represent the breadth of sounds in Spanish) likely contributes to the disparity between the two modalities.

In stark contrast to the low-proficiency group, high-proficiency L2 learners showed no significant ROIs in the auditory modality when comparing with L1 speakers, while still exhibiting one significant ROI in the visual modality. This suggests that stronger nativization processes take place in the auditory modality when improving proficiency from a beginner level to an intermediate level. But minding the difference between correlation and causation, it may also be that general proficiency improves with the consumption of popular Spanish media, such as songs and TV shows, which is often consumed in the auditory modality. Thus, higher-proficiency L2 learners may simply be more comfortable in the auditory modality than low-proficiency learners because of their voluntary consumption of Spanish media.

The question of modality-specific nativization, however, is confounded by familiarity with the Latin script (from English classes), unfamiliarity with native Spanish pronunciation (particularly in contrast to the likely inaccurate pronunciation used during subvocalization), the type of instruction received by participants (probably favoring written materials), and the type and frequency of voluntary exposure to native Spanish media. Coupling these confounding variables with the limited number of participants, limited range of proficiencies among participants, and the effects of inter-participant variability, these L2 results should not be taken as conclusive, but rather as a springboard for further investigation.

### 4.3 Key Findings & Conclusion

In brief summary, with native speakers, we have shown that the N400 is amodal while the P600 has modality-specific features, namely a more gradual slope and a lager peak in the auditory modality. We also observed a negativity over occipital electrodes at 600–700 ms only in the visual modality and a negativity over left temporal electrodes at 1100–1200 ms only in the auditory modality. These two modality-specific voltage distributions align with the primary visual cortex and the primary auditory cortex, respectively, possibly suggesting their involvement in modality-specific language processing streams.

With the L2 learner groups, we have shown greater nativization in high proficiency L2 learners than in low proficiency L2 learners. We also show that low proficiency L2 learners showed less nativelike ERPs in the auditory modality than in the visual modality, where high proficiency L2 learners showed more nativelike ERPs in the auditory modality than in the visual modality. We additionally observed more modality-specific differences in high proficiency L2 learners, suggesting that modality-specific specialization in language processing is a neurological strategy acquired with higher proficiency.

### 4.4 Limitations

Due to geographic constraints, our L1 experiment was limited by a somewhat small pool of participants. Additionally, the stimuli materials used were very easy for native speakers, leading to a ceiling effect that obfuscated any modality effects that may have otherwise manifested in the behavioral data. Furthermore, since materials were compiled from a small list of Spanish words learned in first-semester Spanish class at the participants’ university, features such as word length and stressed syllable could not be controlled for. Although not ideal, we do not believe this would fundamentally change our findings.

Due to the relative lack of interest in Spanish among Japanese university students, our L2 participant pool was also small, and smaller still after dividing into high and low proficiency groups. It is also important to bear in mind that our participants roughly ranged in proficiency from A1 to B1 on the CEFR scale, thus only representing the lower half of possible L2 proficiencies.

As would almost always be the case, there are several confounding variables in L2 acquisition (e.g., familiarity with native pronunciation, focusing studies on written vs spoken materials, etc.) that must not be forgotten when interpreting results.

Finally, since no such studies had been carried out before, the whole epoch and whole scalp distribution had to be analyzed, increasing the likelihood of Type I family-wise errors. On the other hand, we followed the standard Holm-Bonferroni p value correction procedure and tried to be as clear as possible when referring to adjusted or unadjusted p values. Still, it is more statistically robust to approach such phenomena with a specific ROI and time window in mind prior to conducting the experiment. To that end, we hope this study can be used to inform future probing into the neurophysiological modality effect in language processing and encourage all neurolinguistics researchers to make a concerted effort to adequately mind the effect of stimulus modality both while designing experiments and while interpreting results.

